# Neutrophil myeloperoxidase as a functional biomarker for RSV severity: implications for *in vitro* therapeutic screening

**DOI:** 10.1101/2025.10.13.682115

**Authors:** Machaela Palor, Tereza Masonou, Elisabeth J. Robinson, Wenqing Chen, Samuel Ellis, Laura Buggiotti, Amy I. Jacobs, Thomas Benoist, Paolo De Coppi, Jennifer L. Rohn, Gabriele Pollara, Mario Cortina-Borja, Maximillian N.J. Woodall, Robert E. Hynds, Rosalind L. Smyth, Samiran Ray, Claire M. Smith

**Author notes:** Corresponding author: Dr Claire Mary Smith, UCL Great Ormond Street Institute of Child Health, 30 Guilford St, London WC1N 1EH.

## Abstract

Respiratory syncytial virus (RSV) is a leading cause of severe lower respiratory tract infections in infants, yet effective therapeutics are lacking. The aim of this study is to develop an *in vitro* model that recapitulates key clinical outcomes in infants with RSV bronchiolitis to help accelerate the discovery of effective therapeutics.

Neutrophil activation and influx into the airways are hallmarks of severe RSV infection, but these responses are difficult to quantify in clinical trials. Here we profile peripheral blood-derived neutrophils from infants with RSV admitted to the Paediatric Intensive Care Unit (PICU) and identify myeloperoxidase (MPO) as a key indicator of disease severity when compared to age matched controls.

To mechanistically model this response, we established a paediatric airway epithelial air–liquid interface (ALI) system incorporating an endothelial layer and primary neutrophils to recapitulate the tissue microenvironment at the blood-airway barrier. Following RSV infection, neutrophil migration and activation were assessed using flow cytometry. The inclusion of an endothelial layer enhanced physiological relevance and more accurately replicated *in vivo* MPO responses.

We then evaluated two antiviral candidates (remdesivir (RDV) and RSV604) to assess their ability to modulate neutrophil activation. While both compounds reduced viral load at 24 hours post-infection, only RSV604 attenuated MPO expression.

These findings establish MPO as both a biomarker of RSV disease severity and a functional readout of therapeutic efficacy and demonstrate that targeting neutrophil⍰driven inflammatory pathways may be critical for reducing pathology in infant RSV infection.

**Take home message:** Antiviral drug discovery should include neutrophil MPO reduction as a readout of therapeutic efficacy.

**Graphical Abstract:** 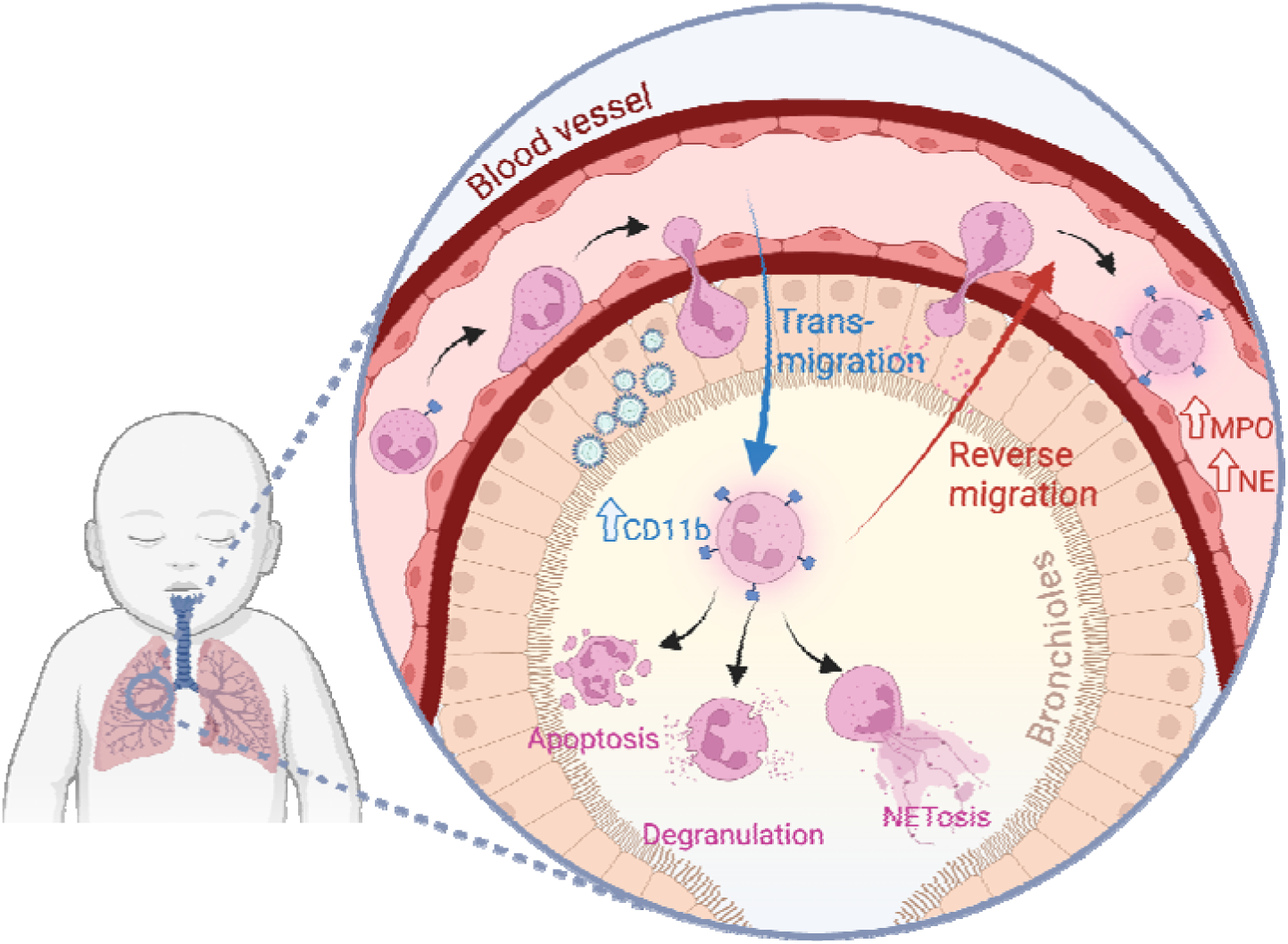

## Introduction

Respiratory syncytial virus (RSV) is a leading cause of severe lower respiratory tract infections in infants and young children, resulting in approximately 3.2 million hospital admissions and an estimated 118,000 deaths in children under five years of age (1). Despite these substantial health impacts, current therapeutic options primarily focus on prevention and symptom management. There are currently no routinely used, effective antivirals for immunocompetent children or adults, and ribavirin, historically the only licensed antiviral, is now rarely used even in immunocompromised populations due to high cost, and concerns about toxicity (2, 3). Prophylactic strategies for high-risk infants include palivizumab, a monoclonal antibody requiring monthly dosing (4, 5) and nirsevimab, a recently approved monoclonal antibody shown to provide extended protection with a single dose (6, 7). The lack of accessible, effective therapies highlights the need for continued advancements in RSV treatment and prevention.

Neutrophils are rapidly recruited to the lungs during RSV infection, playing a key role in the host defence mechanism by targeting the virus (8). However, excessive neutrophil infiltration and activation can contribute to airway inflammation, epithelial damage, and disease severity (9, 10). While mouse studies have shown that neutrophil activation primarily occurs within the lung environment during RSV infection (11, 12), others have shown that neutrophils in the systemic circulation of RSV-infected children also exhibit high levels of activation markers, indicating that their activation is not limited to the lung environment (13, 14). Additionally, RSV proteins have been detected within these circulating neutrophils (15), raising the possibility that some neutrophils, after interacting with virus-infected cells in the lungs, may re-enter the bloodstream. This process of reverse migration could have significant implications for systemic inflammation and viral dissemination.

Understanding neutrophil behaviour and activation during RSV infection, including during their migration across the airway epithelial barrier, is crucial for developing therapeutic strategies to mitigate pathological inflammation without compromising the antiviral response. Studies have shown that trans-epithelial migration is essential for neutrophil activation during RSV infection, leading to higher expression of activation markers and release of inflammatory mediators (11, 12, 16). *In vivo* studies are limited by their complexity and ethical considerations, necessitating the use of *in vitro* models that closely replicate the human airway environment, which can differ between children and adults (17). Differentiated airway epithelial cells (AECs) cultured at the air-liquid interface (ALI) provide a physiologically relevant platform to study neutrophil migration and the effect of antiviral treatments on this process. This approach is particularly valuable given the urgent need for predictive pre-clinical models that can accelerate the development and optimisation of antiviral therapies.

In this study, we aimed to develop an *in vitro* model that recapitulates key clinical outcomes of infants with RSV bronchiolitis. We analysed neutrophils from RSV-infected infants and validated our clinical observations in a human trans-epithelial ALI migration model (11, 18). To enhance physiological conditions, we incorporated a vascular endothelial cell (EC) layer alongside differentiated AECs, enabling us to study early infection dynamics and neutrophil phenotype in a more representative airway environment. Finally, we evaluated the effects of two repurposed antiviral therapies, remdesivir (RDV) (a broad-spectrum nucleoside analogue) and RSV604 (a small-molecule RSV fusion inhibitor), on neutrophil migration, priming, and degranulation, to assess their potential in modifying immune responses during RSV infection.

## Methods

### Neutrophil Isolation and Purification

Ethical approval was obtained from the Living Airway Biobank (19/NW/0171; UCL_CS_23A) for paediatric patients admitted to the GOSH PICU with an indwelling arterial or a central venous line, or UCL Research Ethics Committee (Project ID number: 19165/001) for the collection of venous blood from healthy adult volunteers. Written informed consent was obtained directly from donors, or their guardian and blood sampling was carried out by trained phlebotomists.

Neutrophils were isolated from children in PICU with RSV infection (n = 5, age range 1.5–29 months, 60% female) and uninfected control PICU patients (who have had elective procedures) (n = 5, age range 0.2–7 months, 80% female) paediatric patients (**Fig. 1b** and **Fig. 2b**; detailed patient characteristics in **Table 1**) or healthy adults (n = 15, aged 20-40 years, 58% female). Neutrophils were isolated using the EasySep™ Direct Human Neutrophil Isolation Kit (#19666, STEMCELL Technologies, UK), yielding approximately 1 x 10 neutrophils with up to 99% purity. Briefly, the sample was incubated with 500 µl of the Isolation Cocktail and RapidSpheres™, followed by a magnetic separation in 50 ml of cell separation buffer. The clear fraction was collected, re-incubated with RapidSpheres™, and placed on a magnetic stand. After further incubation and centrifugation at 300 x g, the pellet was resuspended in separation buffer and further enriched using the EasySep™ Human Neutrophil Isolation Kit (#17957, STEMCELL Technologies, UK) as per the manufacturer’s instructions.

**Figure 1.**
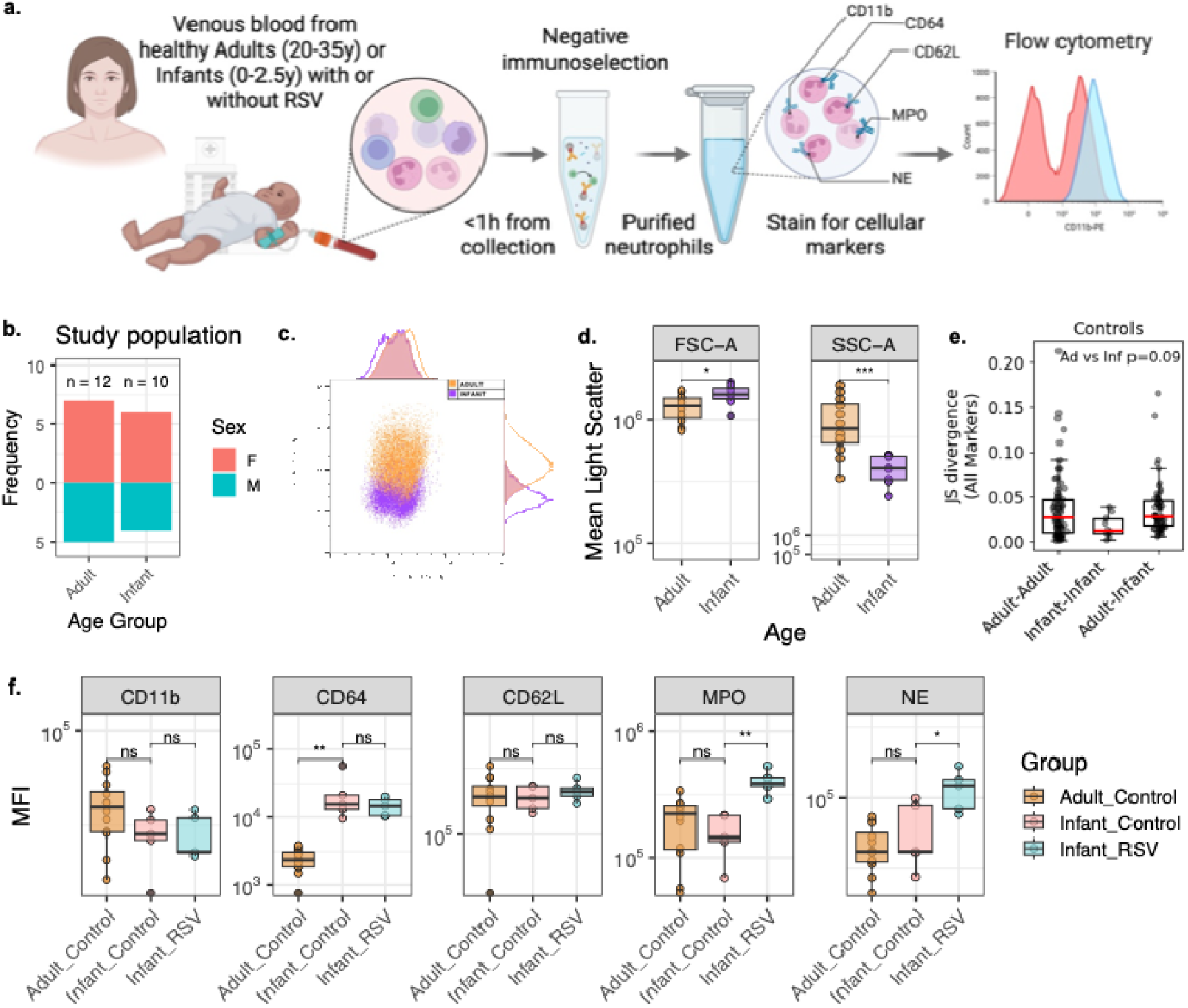
Phenotypic comparison of neutrophils from adults and infants with or without RSV infection. (a) Schematic of study design showing blood sampling from adults and infants with RSV bronchiolitis (RSV) and uninfected control ICU patients (DC). (b) Study population demographics by age group and sex (F = female, M = male). (b) Composition of the study population, showing numbers of adult and infant donors and their sex distribution. (c) Representative flow/JIcytometry plots illustrating neutrophil gating and marker expression profiles. (d) Comparison of forward scatter (FSC/JIA) and side scatter (SSC/JIA) between adult and infant neutrophils, indicating age/JIassociated differences in cell size and granularity. (e) Jensen–Shannon divergence (JSD) analysis of control donor by age. (f) Median fluorescence intensity (MFI) of neutrophil activation and granule markers (CD11b, CD64, CD62L, MPO, NE). Boxplots display median and interquartile range (n=5-10). Significance levels are indicated.

**Figure 2.**
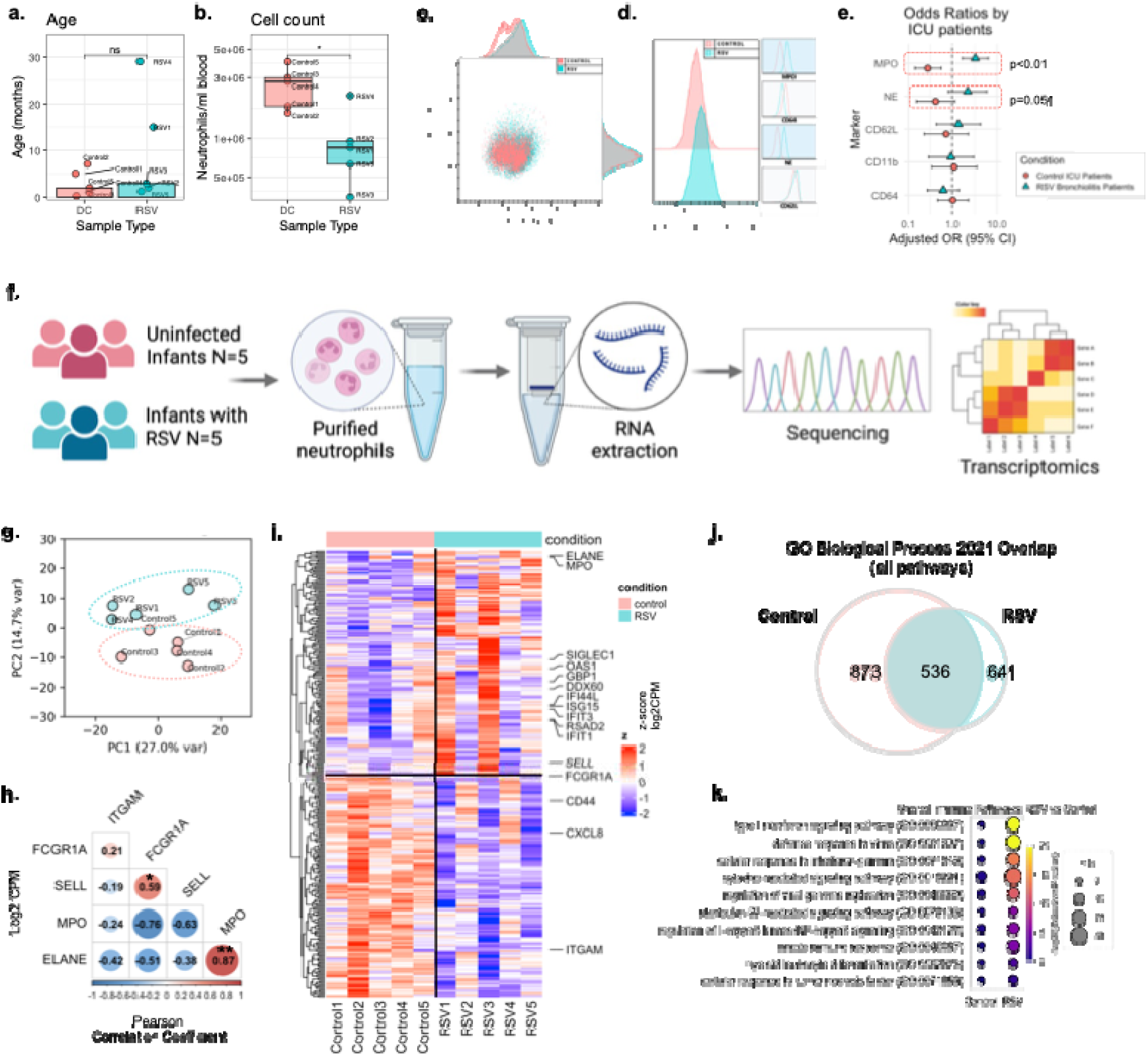
Neutrophil phenotyping and transcriptomic profiling in RSVh infected infants **(a)** Age distribution of RSV/JIinfected infants and uninfected controls included in the study. **(b)** Absolute neutrophil counts in peripheral blood from control and RSV/JIinfected infants. **(c–d)** Flow/JIcytometric analysis of circulating neutrophils. **(c)** Representative scatter plots illustrating the neutrophil gating strategy. **(d)** Representative histograms showing expression levels of key neutrophil surface markers. **(e)** Odds ratio plot showing relative likelihood of marker expression changes in RSV versus control patients; points represent odds ratios with 95% confidence intervals. **(f)** Schematic illustrating sample processing workflow, including neutrophil purification, RNA extraction, and transcriptomic analysis. Made using BioRender.com. **(g)** Principal component analysis (PCA) of neutrophil transcriptomes from RSV/JIinfected and control infants, demonstrating clear separation by PC2 based on infection status. **(h)** Correlation matrix of transcript levels of key neutrophil marker genes. Error bars represent standard deviation; ns = not significant; *p < 0.05, **p < 0.01. **(i)** Heatmap of differentially expressed genes (DEGs) between RSV and control neutrophils. Rows represent individual genes, columns represent donors, and colours indicate z/JIscore–scaled expression. Highlighted genes denote strongly up- or down/JIregulated transcripts. **(j)** Venn Diagram of gene ontology (GO) enrichment analysis of DEGs, showing number of pathways associated with Control or RSV neutrophils. (**k**) Dot/JIplot illustrating the top enriched GO Biological Process pathways derived from differentially expressed genes. Dot size represents the number of genes contributing to each pathway, and dot colour indicates adjusted p/JIvalue. Pathways are grouped according to enrichment in RSV/JIinfected or control neutrophils.

**Table 1.**
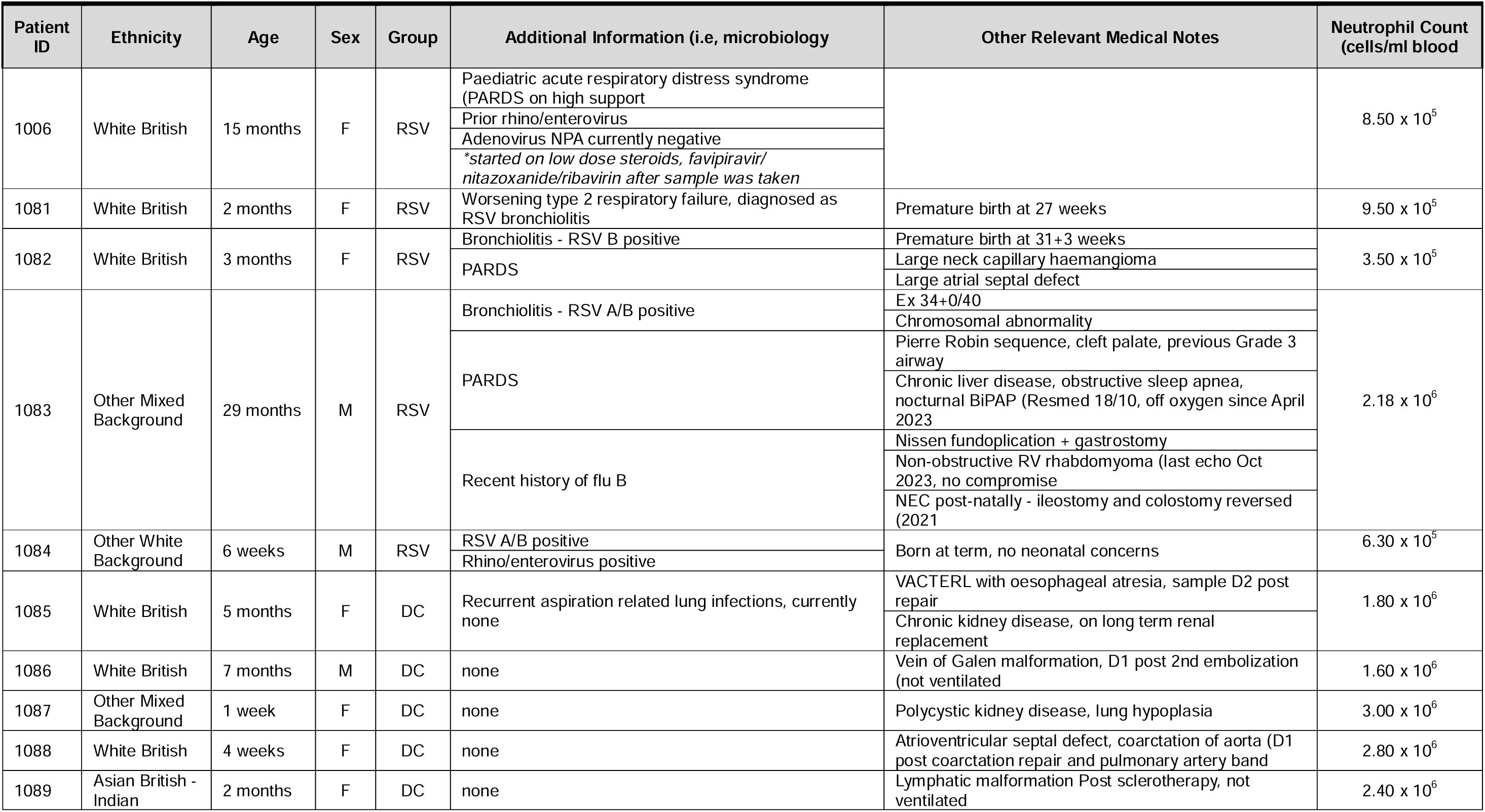
PICU Patient Demographics.

**Table 1b.** Adult Demographics.

| Patient ID | Ethnicity | Age (years) | Sex | Group |
| --- | --- | --- | --- | --- |
| 930 | Asian | 25.16 | F | Healthy Adult |
| 961 | White British/Irish | 23.52 | F | Healthy Adult |
| 964 | White British/Irish | 22.55 | F | Healthy Adult |
| 1000 | White British/Irish | 24.95 | M | Healthy Adult |
| 1001 | Asian | 27.54 | F | Healthy Adult |
| 1004 | White British/Irish | 24.16 | F | Healthy Adult |
| 1005 | Asian | 24.61 | M | Healthy Adult |
| 1008 | White British/Irish | 22.39 | F | Healthy Adult |
| 1077 | White British/Irish | 31.14 | M | Healthy Adult |
| 1078 | White British/Irish | 34.40 | M | Healthy Adult |
| 1079 | Asian | 29.22 | M | Healthy Adult |
| 1080 | White British/Irish | 27.67 | F | Healthy Adult |

### Airway Epithelial Cell Culture and Differentiation

Primary airway epithelial cells (AECs) were obtained from adults (n=6, 19-40y as described in (12)) and children (n=5, 0-4y) by nasal brushings. Primary paediatric (2-year-old female Caucasian) bronchial AECs were purchased from Epithelix Sàrl (Geneva, Switzerland) and co-cultured on mitotically inactivated 3T3-J2 feeder cells as described previously (19, 20). Cell stocks were frozen down at 1 x 10^6^ cells/100 µl in Bambanker™ BB01 (NIPPON Genetics, Germany), a serum-free cryopreservation medium.

AECs were then cultured at ALI in complete PneumaCult™-ALI medium (STEMCELL Technologies, Vancouver, Canada) for 28 days to promote differentiation as described previously (21).

### Endothelial Cell Culture

Human umbilical vein endothelial cells (HUVECs) transduced with ETS variant transcription factor 2, also known as ‘reset’ vascular ECs (22), were cultured on recombinant laminin-511 E8 fragment-coated flasks (iMatrix-511, AMSBIO, UK) in endothelial growth medium composed of complete Endopan 300 SL medium (Pan BioTech, Germany) and 5% human serum (Merck, Darmstadt, Germany). These reagents proved to be the most effective growth conditions for endothelial cells of others tested (**Supplementary Fig. 2**). Cell stocks were frozen down at 1 x 10^6^ cells/ml in EGM supplemented with 40% HS and 10% DMSO.

For co-culture at ALI, ECs were seeded into membrane inserts at 3.5 x 10^4^ in 100 μl complete PneumaCult™-ALI medium supplemented with iMatrix-511 (1.5 μg/cm^2^) and incubated for 24 hours prior to performing dextran permeability assays or neutrophil migration experiments.

### Dextran Permeability Assay

To assess cell barrier integrity, membrane inserts with ALI AEC cultures alone or in co-culture with ECs, were placed in Hanks’ balanced salt solution containing calcium and magnesium (HBSS+/+) plus Texas Red™-dextran (Thermo Fisher Scientific, UK) at 100 µg/ml for 20 minutes in the dark. 100 µl of standards and supernatant were transferred to a solid black 96-well plate in triplicate and fluorescence was measured with a BMG FLUOstar OMEGA microplate reader at an excitation/emission wavelength of 595/615 nm. A membrane without cells served as a control. The translocated dextran concentration, derived from a standard curve, indicated paracellular permeability.

### Viral Infection and Quantification

Recombinant GFP tagged-RSV A2 strain was provided by Fix et al (23). ALI cultures were infected at a MOI of 1 in 25 µl HBSS+/+ for one hour at 37°C. The virus quantity was based on an estimated 1 x 10^5^ cells per membrane. Mock-infected controls received 25 µl HBSS+/+. After inoculation, the virus solution was removed, and cultures were incubated for 24 hours with replenished basolateral ALI medium.

For quantification, viral RNA was extracted from apical supernatants using the QIAamp Viral RNA Mini Kit (Qiagen, Germany). cDNA was synthesized from 100 ng RNA using the High-Capacity RNA-to-cDNA Kit (Thermo Fisher Scientific, UK) in a 40 μl reaction. RT-qPCR was performed using TaqMan Universal Master Mix II (Thermo Fisher Scientific, UK) with specific primers and a fluorescent probe for the RSV-A N protein as described previously (21). Samples were run on an AB Biosystems StepOnePlus Real-Time PCR System (Thermo Fisher Scientific, UK). Viral load was extrapolated from a standard curve generated by performing ten-fold serial dilutions of a plasmid containing the N protein sequence starting at 1 x 10^6^ copies to 1 copy (24).

### Trans-epithelial Migration Assay

On the day of the neutrophil migration assay, ALI cultures were incubated with 600 µl HBSS+/+ for one hour at 37°C to collect immunomodulatory factors. Supernatants were collected, centrifuged, and 400 µl was transferred to a 24-well plate with ALI cultures. Purified neutrophils were diluted to 5 x 10^6^ cells/ml in HBSS+/+ with 1% (v/v) autologous human serum. 100 μl was added basolaterally to each membrane insert and incubated for one hour at 37°C.

### Assessment of Neutrophil Markers and Degranulation

After migration, neutrophils (apical, adherent, basolateral) were collected, centrifuged, blocked with Human TruStain FcX™, and stained with LIVE/DEAD™ fixable violet dye (Thermo Fisher Scientific, UK). Following washing, cells were stained with 1/50 dilution of CD11b-FITC (Miltenyi Biotec, 130-110-552), CD64-APC-Cy7 (Miltenyi Biotec, 130-116-199), and CD62L-PE-Cy7 (Miltenyi Biotec, 130-129-810) for 20 minutes at 4°C in the dark. Cells were washed, fixed with PFA, permeabilised, and stained intracellularly with MPO-APC (Miltenyi Biotec, 130-119-786) and NE-PE (Santa Cruz Biotechnology, sc-55549 PE). After a final wash, neutrophils were resuspended in FACS buffer and analysed using a CytoFLEX S flow cytometer, obtaining mean fluorescence intensity (MFI) values for CD11b, CD64, CD62L, MPO, and NE from the CD11b-positive gated population. The gating strategy for the identification of neutrophils by flow cytometry is shown in **Supplementary Fig. 1**. Data are presented as raw MFI values, or to control for day⍰to⍰day variability, we normalised MFI from migration experiments to same⍰day “medium⍰only” controls and use FMO and cytometer calibration/PMT stability checks. To visualize patterns of marker expression, we utilized a Quartile Score, where the mean MFI of each marker was categorized into quartiles (0–3); 0 corresponds to the lowest quartile and 3 to the highest, calculated across all compartments and experimental conditions. Associations between marker expression patterns across different compartments or conditions of the model were visualised using `radarcharts` generated with the `fmsb` R package (25). Here, the centre of each plot represents the lowest level of marker expression, while points closer to the edges indicate higher expression levels. The individual quartile scores were totalled to calculate a Combined Quartile Score for each condition. Statistical analyses were performed using Wilcoxon signed-rank tests, with adjustments for multiple comparisons to control the false discovery rate.

Neutrophil elastase activity in supernatants was measured using the fluorometric Neutrophil Elastase Activity Assay Kit (Item No. 600610, Cayman Chemical, Michigan, USA) as per the manufacturer’s instructions. Samples were diluted 1/10, incubated with elastase substrate, and fluorescence was measured at 485/525 nm with a BMG FLUOstar OMEGA microplate reader.

For MPO activity, samples were diluted 1/2 and incubated with 3,3′,5,5′-Tetramethylbenzidine substrate as described using the colorimetric Neutrophil MPO Activity Assay Kit (Item No. 600620 Cayman Chemical, Michigan, USA). Absorbance was measured at 650 nm with a BMG FLUOstar OMEGA microplate reader.

### Transcriptomics

Neutrophil RNA from RSV-infected and control paediatric patients was extracted using the RNeasy Plus Mini Kit (Qiagen, Germany). RNA samples were then reverse-transcribed and sequenced using the cDNA-PCR Sequencing V14 - Barcoding kit (SQK-PCB114.24) according to the manufacturer’s instructions. Samples were sequenced on the R10.4.1 flow cell with a MinION Mk 1C sequencer (Oxford Nanopore Technologies, UK). Raw sequencing data were base-called during acquisition via MinKNOW (Oxford Nanopore Technologies). Reads were processed using the ONT EPI2ME: wf-transcriptomes workflow to generate aligned BAM files against the human reference genome GRCh38.p14.115. Gene-level counts were subsequently quantified from the aligned BAM files using featureCounts (Subread package), generating a raw gene count matrix for downstream analysis. Differential gene expression between RSV and control groups was conducted using the edgeR package. Lowly expresses genes were filtered by retaining genes with CPM ≥ 1 in at least two samples. Library size normalization was performed using the trimmed mean of M values (TMM) method implemented in edgeR. Principal component (PC) analysis was performed on log2-transformed normalized expression values (log2(CPM + 1) using the top 2000 most variable genes. To visualise genes contributing to PC2 separation, the top 150 genes with the highest positive and 150 genes with the highest negative PC2 loadings were selected corresponding to RSV-enriched and Control-enriched gene sets, respectively scaled (z-scored per gene) and visualised using hierarchical clustering heatmaps. Gene ontology (GO) enrichment analysis was performed using Enrichr through GSEApy Python interface, focusing on GO Biological Process 2021 annotations. Enrichment significance was evaluated using adjusted P-values (Benjamini-Hochberg correction) and enrichment strength was quantified using the Enrichr combined score. Shared GO terms between RSV-up and Control-up gene sets were identified.

### Cytokine Measurement

The BD Cytometric Bead Array (CBA) assay (BD Biosciences, USA) was used to quantify human soluble proteins, including key mediators of inflammation and neutrophil migration, IL-6, CXCL8 (IL-8), and CXCL10 (IP-10) (26, 27), in supernatants from mock- or RSV-infected cultures. Following the manufacturer’s protocol, a standard curve was generated, and test samples were diluted 1/10 in assay diluent. Standards and samples were incubated with capture beads for one hour, followed by PE detection reagent for two hours. After washing, the beads were resuspended in wash buffer and analysed using a BD FACSymphony A5 flow cytometer (BD Biosciences, Franklin Lakes, NJ, USA).

### Antiviral Treatment

RSV-infected bronchial AECs were treated with a broad-spectrum antiviral remdesivir (RDV, Bio-Techne, USA) or RSV-specific antiviral compound, RSV604 (Cambridge Bioscience, UK), which has high efficacy in inhibiting RSV replication (28). RDV (Bio-Techne, USA) and RSV604 (Cambridge Bioscience, UK) were diluted in ALI medium to working concentrations of 6.6 μM and 10 μM, respectively, and added to the basolateral side at four hours post-infection. These concentrations were based on RDV EC50 values (29) and previously reported non⍰cytotoxic RSV604 doses effective against RSV in human AEC (28). AECs were used for neutrophil migration experiments at 1 day post infection (dpi). The effects of the antiviral on RSV-induced neutrophil migration and receptor expression were assessed by comparing treated and untreated cultures. Additional infected AECs with and without antiviral treatment were recorded for viral GFP fluorescence using a Nikon Ti-E microscope up to 7 dpi.

### Statistical Analyses

Data were analysed and visualized using GraphPad Prism v10.0 or RStudio 2023.09.1+494. It is important to note that there was considerable variability (>1 log) in baseline neutrophil marker expression across different donors (**Supplementary Fig. 3**). This variability likely reflects inherent differences in neutrophil states between individuals at collection. To account for this in our analysis, we present our data as the mean fold change in MFI relative to neutrophils cultured only in medium (‘medium-only’) for each condition. This approach normalized the individual baseline differences and allowed us to focus on the migration-dependent effects.

Normality was assessed with the Shapiro-Wilk test. For normally distributed data, a student’s t-test (two groups) or One-way ANOVA with Bonferroni correction (three or more groups) was used. Two-way ANOVA with Tukey correction was applied for comparisons involving two factors. Non-normally distributed data were analysed with Mann-Whitney U or Wilcoxon signed-rank tests (two groups) and Kruskal-Wallis or Friedman tests with Dunn’s post-hoc for unpaired or paired data, respectively. Results are presented as mean ± SEM unless specified otherwise. Statistical significance was set at *P* < 0.05.

For age-adjusted analyses, data were log⍰transformed and analysed in R using linear and logistic regression to assess effects of RSV status and age, with effect ratios, odds ratios, and predicted values derived using broom, questionr, and ggeffects.

## Results

### Comparable Peripheral Blood Neutrophil Phenotypes Across Age with Higher Neutrophil Degranulation Markers in RSV-Infected Infants

To examine age⍰related differences in neutrophil phenotype and how these may be altered by RSV infection, we analysed peripheral blood neutrophils from healthy adults and infants, including both uninfected controls and RSV⍰infected infants (**Fig. 1a**). Adults and infants were balanced by sex (**Fig. 1b**), with detailed donor characteristics provided in TableD1.

Flow⍰cytometric analysis revealed that infant neutrophils exhibited lower forward scatter (FSC) and side scatter (SSC) compared with adult neutrophils (**Fig. 1c**), indicating reduced cell size and granularity. These differences were consistent across donors (**Fig. 1d**) and align with previously described distinctions between neonatal/infant and adult granulocytes (12). There was no difference in FSC or SSC between uninfected controls and RSV⍰infected infants (**Supplementary Fig. 4a**)

To assess phenotypic similarity across age groups, we calculated the Jensen–Shannon divergence (JSD) using uninfected control donors only. JSD did not differ significantly between adults and infants (pD=D0.09; **Fig. 1e**), indicating that the overall multivariate neutrophil phenotypes were broadly comparable at baseline. In controls, activation and granule marker expression was also largely similar across ages, with CD64 the only marker found to be significantly higher in infants at baseline (**Fig. 1f–g**). Relative to infant controls, age-adjusted analyses showed significantly higher MPO and NE expression (pD<D0.05) in RSV⍰infected infants, whereas CD11b, CD64 and CD62L remained unchanged (**Fig. 1f–g, Supplementary Fig. 4b**).

### RSV Infection Drives Neutrophil Phenotypic and Transcriptional Reprogramming in Infants

We then profiled wider phenotypic changes in neutrophils from RSV⍰infected infants and age⍰matched uninfected controls (Fig. 2a). We found that peripheral blood neutrophil counts were significantly lower in RSV⍰infected infants, with a mean viable cell count of 9.9 × 10D cells/mL compared to 2.6 × 10D cells/mL in control infants (p < 0.05) (Fig. 2b), consistent with enhanced tissue recruitment during infection (8, 30, 31).

Flow cytometric analysis (**Fig. 2c**) showed differences in neutrophil marker expression between the groups (**Fig. 2d**) and odds⍰ratio analysis confirmed that MPO and NE were the markers most strongly enriched in RSV infection, indicating a significantly increased likelihood of elevated expression relative to controls (**Fig. 2e**).

To understand infection⍰driven changes beyond cell protein expression, we performed transcriptomic profiling of purified neutrophils (Fig. 2f). Principal component analysis (PCA) revealed a clear separation between RSV⍰infected and control infants (Fig. 2g), indicating an infection⍰associated transcriptional programme. Gene expression correlation analysis across both conditions showed significant correlation between *ELANE* (encoding NE) and *MPO* (encoding MPO) (Fig. 2h). Differential expression analysis identified RSV⍰responsive genes (Fig. 2i) and gene ontology enrichment revealed 105 pathways uniquely associated with RSV infection and 536 pathways shared between groups (**Fig. 2j**). RSV induced marked up⍰regulation of antiviral, interferon⍰stimulated, cytokine⍰responsive and inflammatory pathways compared to control neutrophils (**Fig. 2k**). Together, these findings highlight a broad transcriptional reprogramming of circulating neutrophils in response to RSV infection.

### RSV Infection Increases Neutrophil Migration and Activation in a Paediatric Airway Epithelium Model

To test whether age-associated interactions with the airway microenvironment drives the heightened neutrophil degranulation observed in blood, we infected well⍰differentiated air–liquid interface (ALI) nasal epithelial cultures from children and adult donors with RSV or mock control (Fig. 3a). After 24h infection, transepithelial electrical resistance (TEER) was not reduced by RSV infection relative to mock, with no loss of barrier integrity in either age group (Fig. 3b).

**Figure 3.**
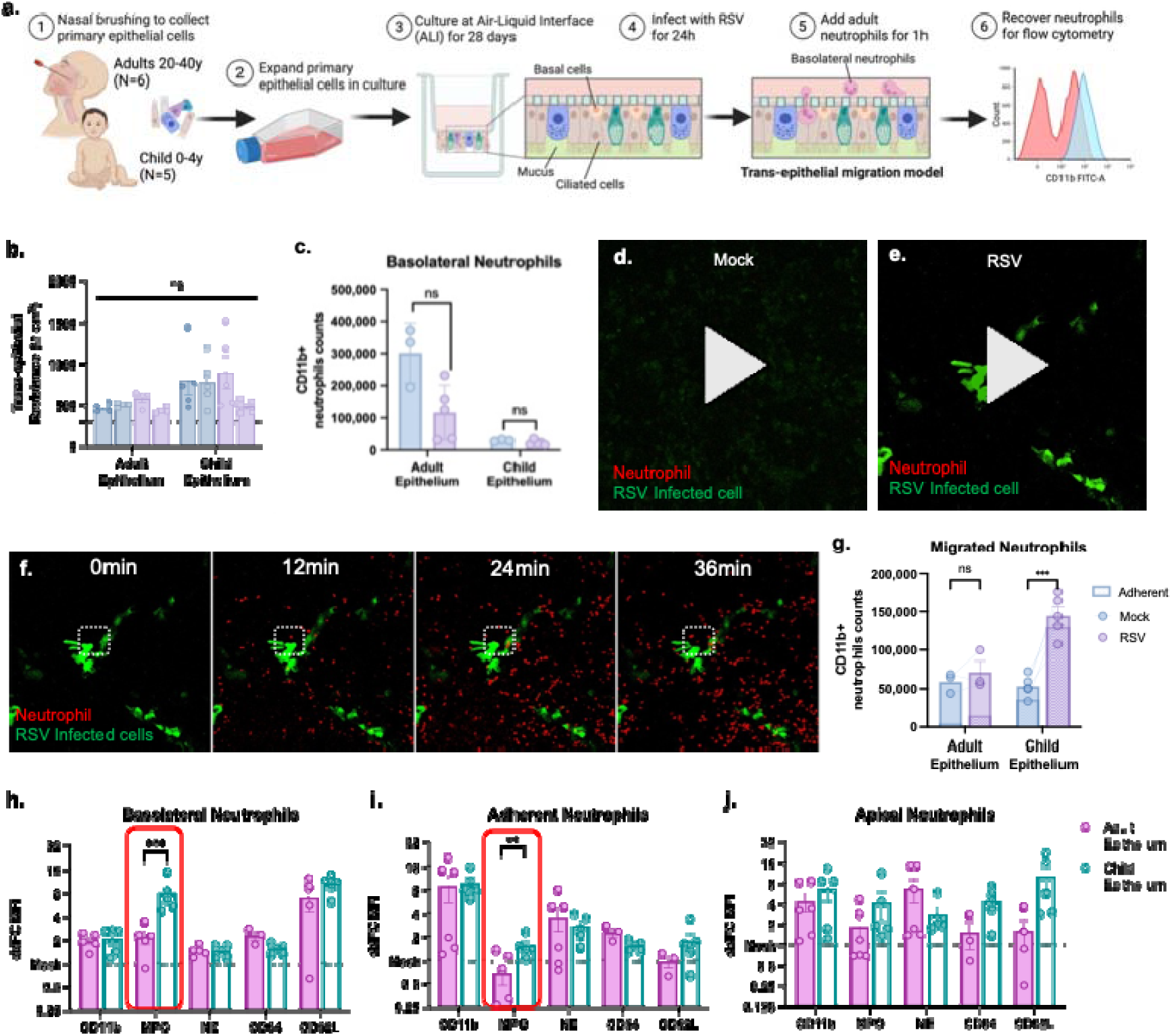
RSV infection enhances neutrophil migration across paediatric airway epithelium and alters neutrophil marker expression. **(a)**, Schematic of experimental workflow. **(b)**, Trans/JIepithelial electrical resistance (TEER) of adult and child ALI cultures under mock or RSV/JIinfected conditions, measured prior to neutrophil assays. Bars show mean ± SEM. ns = not significant. **(c)** Quantification of basolateral neutrophils in adult or child ALI cultures under mock or RSV infection. Mean ± SEM; p < 0.05, p < 0.01, p < 0.001. **(d–e),** Timelapse videos showing neutrophils (red) interacting with (d) mock/JIinfected and (e) RSV/JIinfected epithelial cultures. RSV/JIinfected epithelial cells are shown in green. Imaged every 2 minutes for 1 hour from the addition of neutrophils using an inverted Zeiss LSM 710 confocal microscope. Scale bars, as indicated. **(f)**, Image stills from (d) white boxed area indicates zone showing of loss of GFP-positive cell. **(g)** Quantification of migrated neutrophils following migration across adult or child ALI cultures under mock or RSV infection. Mean ± SEM; p < 0.05, p < 0.01, p < 0.001. **(h–j)**, Flow/JIcytometric characterization of neutrophil activation following migration across adult or child epithelium. Expression of CD11b, MPO, NE, CD64, and CD62L is shown as ΔMFI relative to mock infected controls for neutrophils collected from (h) basolateral, (i) adherent, and (j) apical compartments. Bars show mean ± SEM; ns = not significant, **p < 0.05,*** p < 0.01. Only those with p<0.05 are shown in h-j.

We then applied a trans⍰epithelial migration assay in which purified adult neutrophils were added to basolateral side for 1h. Timelapse fluorescence imaging demonstrated increased accumulation of neutrophils at apical sites of RSV infection (Fig. 3d**-f**). Quantitatively, RSV infection increased neutrophil transmigration, with a greater number migrating across the child epithelium than adult epithelium (Fig. 3g). This suggests that the paediatric airway induces stronger recruitment signals during RSV infection. Notably, basolateral neutrophil numbers did not differ between age groups (Fig. 3c).

We next profiled neutrophil phenotypes after exposure to RSV⍰infected paediatric and adult epithelium by flow cytometry. Here basolateral neutrophils exposed to RSV-infected paediatric epithelium displayed higher expression of the degranulation marker MPO compared to the respective adult. Other markers, such as NE, CD11b, CD64, and CD62L remained largely unchanged between age groups (Fig. 3h). A similar pattern was observed in adherent neutrophils at the epithelial interface (Fig. 3i), with paediatric epithelium consistently displaying higher MPO expression relative to mock and compared to the respective adult epithelium. No differences in neutrophil marker expression were observed in apical neutrophils recovered after full transmigration of the epithelial layer (Fig. 3j). Together these results indicate that RSV⍰infected paediatric epithelium drives both enhanced neutrophil recruitment and selective degranulation, aligning with the higher MPO/NE expression observed in infant blood.

### Endothelial Cells Modulate Cytokine Response and Neutrophil Migration Without Affecting RSV Replication

Next, we aimed to establish a more physiologically relevant airway microenvironment by incorporating a vascular endothelial layer to simulate neutrophil extravasation during RSV infection (Fig. 4a). Initial experiments demonstrated that introducing ECs into AEC cultures significantly decreased dextran permeability across the membrane insert compared to empty wells (Fig. 4b) and preserved TEER values around 200 Ω·cm² (Fig. 4c). These findings suggest that the addition of an EC layer restricts passive permeability between the basolateral and apical compartments. Viral load measurements revealed no significant difference between conditions with or without ECs, indicating that ECs do not influence RSV replication (Fig. 4d). However, the cytokine response, including key mediators of inflammation and neutrophil migration, IL-6, IL-8, and IP-10, was significantly higher in apical supernatants from EC+AEC co-cultures at baseline (mock) compared to cultures lacking ECs. RSV infection further increased the cytokine response from baseline, irrespective of EC co-culture conditions (Fig. 4e).

**Figure 4.**
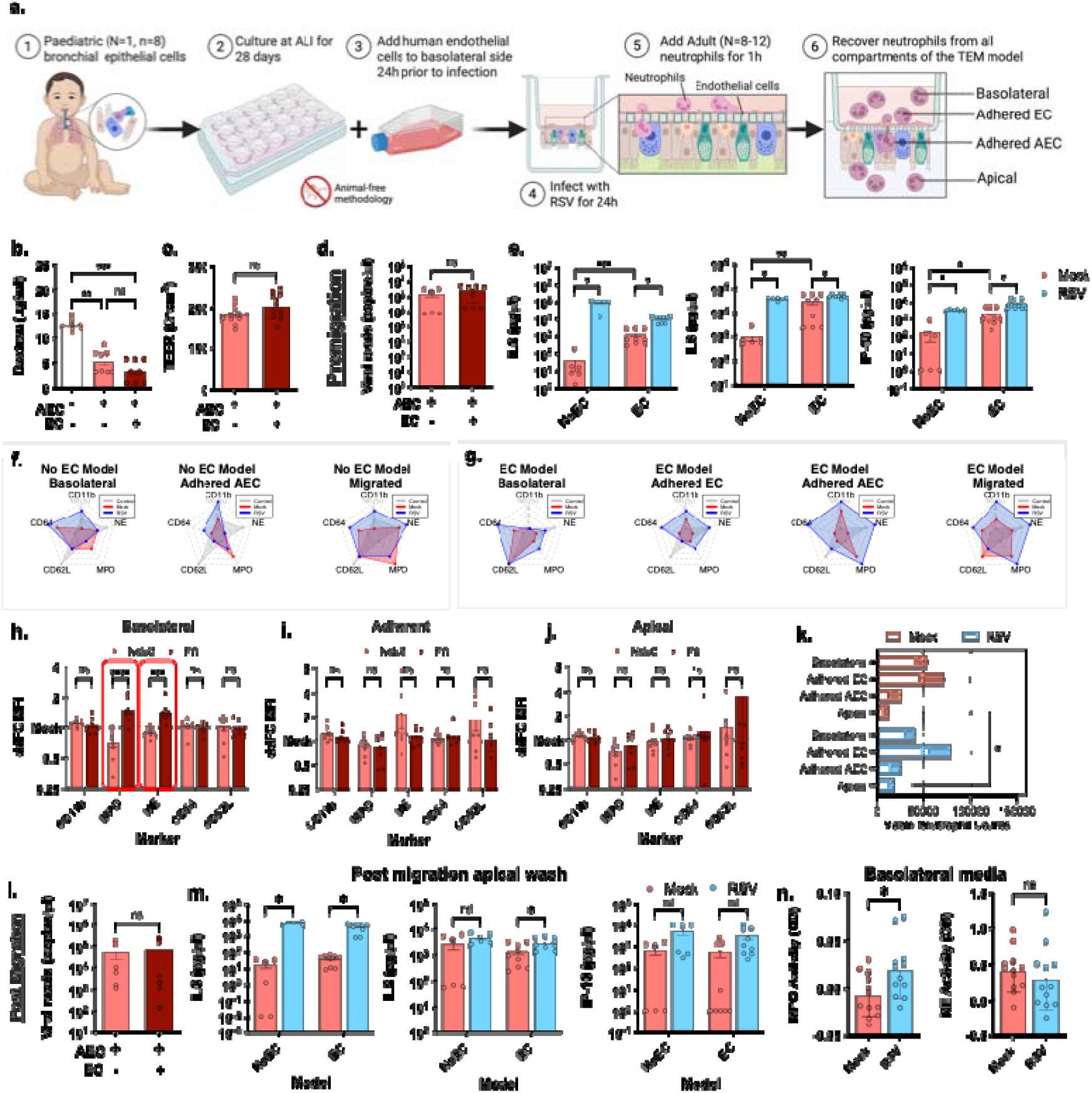
Endothelial–epithelial crosstalk shapes neutrophil responses to RSV infection in a co-culture airway model. **(a)** Schematic of the different co-culture models. Paediatric airway epithelial cells (AECs) were differentiated at ALI for 28 days and combined with human endothelial cells (ECs) on the basolateral side of Transwells. Cultures were infected apically with RSV for 24 h before addition of freshly isolated human neutrophils to the apical chamber. Neutrophils were recovered from the apical wash, adherent epithelial compartment, and basolateral/endothelial compartment for downstream analysis. **(b–e),** Barrier function and cytokine responses of AEC monocultures (AEC) and AEC+EC cocultures (AEC + EC) under mock or RSV conditions. (b) TEER, (c) FITC/JIdextran permeability, (d) Viral reads (e) IL/JI6, IL/JI8 and IP-10 secretion from AEC or AEC + EC cultures under mock or RSV infection. Mean ± SEM. **(f–g)** Radar plots showing quartile scores for neutrophil activation markers after migration through mock/JI or RSV/JIinfected cultures, as determined by flow cytometry. The centre of each plot represents the lowest level of marker expression, while points closer to the edges indicate higher expression levels. **(h-j)**, Paired flow cytometric analyses of neutrophils recovered from NoEC vs. EC models under RSV infection (data shown is ΔΔFC of MFI relative to medium-only and mock controls) across the (h) basolateral, (i) adherent, and (j) apical compartments. Mean ± SEM; **(i)**, Total viable neutrophil counts recovered from each compartment. RSV drives increased neutrophil transmigration to the basolateral and adherent compartments. **(j)**, Viral reads post-migration **(k)**, Neutrophil/JIderived factors in post/JImigration basolateral supernatant from mock/JI or RSV/JIexposed co-cultures: (i) MPO (ng/mL), (ii) IL/JI8 (pg/mL), (iii) IL/JI1β (pg/mL). ns = not significant, *p < 0.05, **p < 0.01, ***p < 0.001*, ****p < 0.0001.

### Endothelial Cells Heighten Degranulation Marker Expression and the Cytokine Response Following Trans-Epithelial Migration During RSV Infection

Analysis of neutrophil markers by flow cytometry indicated different expression patterns of neutrophil markers across different compartments of the model and infection conditions (Fig. 4f**-g**). Across both models (No EC and EC), RSV infection (blue) consistently induces the highest quartile scores for neutrophil marker expression compared to baseline (grey) or mock conditions (red). Paired analysis of neutrophil markers revealed that MPO and NE expression was significantly increased in neutrophils recovered from the basolateral compartment in the presence of ECs (Fig. 4h**-j**, Supplementary Fig. 6a-c). This corresponded with enhanced neutrophil migration, resulting in a greater number of viable neutrophils detaching from the EC+AEC model and migrating to the apical compartment (Fig. 4k**, Supplementary** Fig. 6d).

Viral load measurements post-migration showed no significant differences between non-EC and EC co-culture conditions, indicating that neutrophil transendothelial migration does not influence baseline viral replication dynamics (Fig. 4l, Supplementary Fig. 6e&f). In contrast, cytokine analysis demonstrated significantly higher concentrations of IL-6 and IL-8 following RSV infection (Fig. 4m). Supporting these findings, we also detected significantly higher MPO activity in the basolateral supernatant during RSV infection compared to mock controls (Fig. 4n). Together, these data suggest that endothelial⍰mediated priming of neutrophils enhances their degranulation potential and contributes to an amplified inflammatory response following trans⍰epithelial migration during RSV infection.

### Higher Basolateral MPO and NE in Response to RSV Infection is Dependent on Direct Contact with RSV-Infected AECs

To investigate whether the higher expression of MPO on basolateral neutrophils was driven by soluble factors from ECs or RSV-infected AECs, and was independent of neutrophil trans-epithelial migration, AECs were cultured on membranes with either permissive (3.0 µm pores) or blocked (0.4 µm pores) inserts (Fig. 5a). The smaller pores prevent neutrophil migration while still permitting passive diffusion of soluble factors.

**Figure 5.**
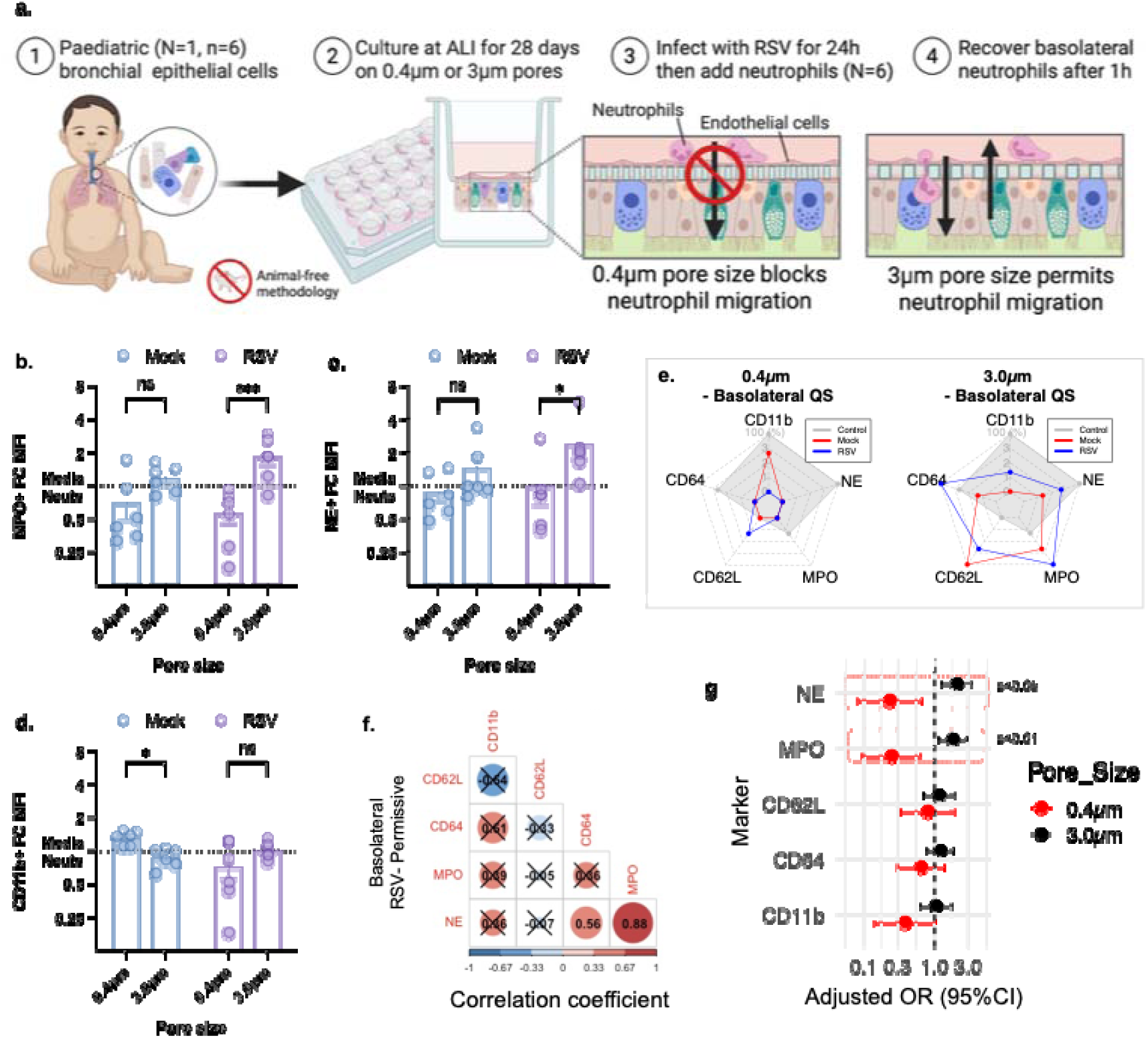
Neutrophil marker expression in response to RSV Infection in the basolateral compartment when neutrophils are permitted to migrate, or migration is blocked. (a) Schematic (b-d) ΔΔFold change (FC) in MFI for individual markers (MPO, NE. CD11b)) on neutrophils recovered from basolateral side under permissive (3.0μm) and blocked (0.4μm) conditions (*p < 0.05, **p < 0.01). (e) Radar plots showing quartile scores (QS) of neutrophil activation markers (CD11b, CD62L, CD64, MPO, NE) under permissive (3.0μm) and blocked (0.4μm) conditions (*p < 0.05 compared to mock). (f) Correlation matrix of marker expression under permissive RSV conditions. Positive (red) and negative (blue) correlations are displayed; strength is indicated by circle size and colour intensity. X = non-significant correlations. (g) Odds ratio plot showing the relative likelihood of marker expression changes under RSV vs. mock conditions in the basolateral compartment with 0.4μm or 3.0μm pore sizes. Points indicate odds ratios with 95% confidence intervals. Red boxes indicate p<0.05.

Using this system, basolateral neutrophils exhibited significantly higher MPO (Fig. 5b**, Supplementary Fig.7a**) and NE expression (Fig. 5c) in RSV-infected cultures under permissive (3.0 µm) conditions compared to blocked (0.4 µm) conditions (P < 0.05). No change in CD11b expression was detected with RSV infection (Fig. 5d) Radar plot analysis further showed that neutrophils in the blocked (0.4 µm) model exhibited lower activation scores across all markers compared to permissive cultures (Fig. 5e).

A correlation analysis demonstrated significant positive associations between MPO and NE in basolateral neutrophils following migration across RSV-infected cultures under permissive (3.0 µm) conditions (Fig. 5f**, Supplementary Fig.7b&c**), suggesting co-expression of these markers. These findings suggest that increased MPO and NE in basolateral neutrophils is dependent on direct contact with RSV-infected AECs, rather than solely due to soluble infection-related factors. The odds ratio analysis (Fig. 5g) supports this interpretation, revealing that MPO and NE expression is dependent on neutrophil migration and shows a marked separation of odds ratios between pore sizes.

### Antiviral Treatments Reduce Viral Load but Not Inflammatory Cytokine Production

Next, we investigated the effects of two existing antiviral agents, remdesivir (RDV) and RSV604, on neutrophil-mediated inflammation (timeline shown in Fig. 6a**&b**), focusing on the production of IL-6, IL-8, and IP-10 and neutrophil marker expression.

**Figure 6.**
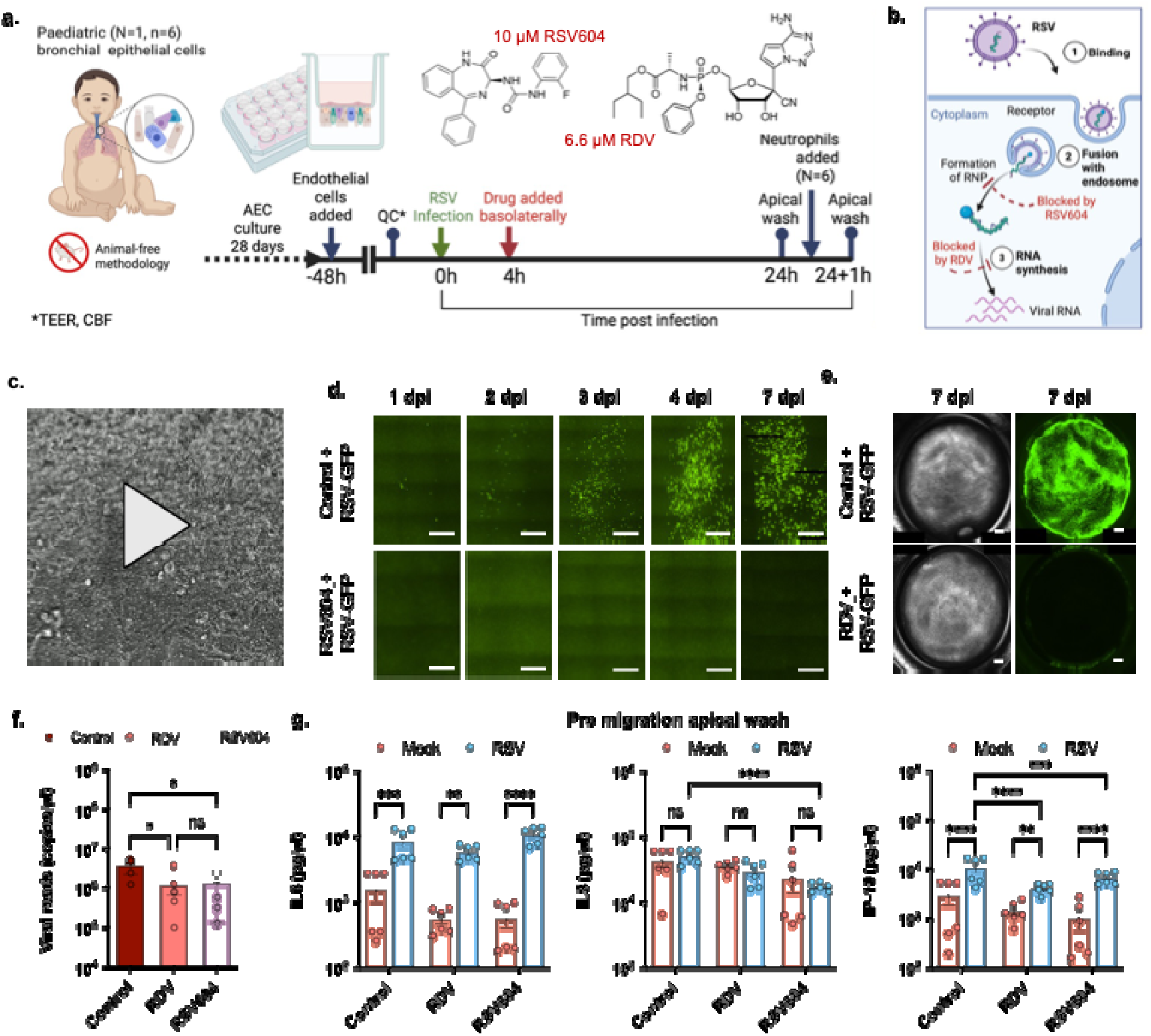
Effects of Antiviral Treatments on viral load and inflammation using in vitro RSV Models. **(a)** Schematic of experimental design showing epithelial infection with RSV, antiviral treatments (RSV604, RDV), and neutrophil migration. Created with BioRender.com. Chemical structures of RSV604 and RDV also shown. (b) Proposed mechanisms of action of RSV604 and RDV during the RSV life cycle. (c) High-speed video of motile cilia demonstrating epithelial function. (d) Representative fluorescence micrographs showing inhibition of RSV-GFP infection in AECs up to 7 days post-infection (dpi) following treatment with 10 μM RSV604. Scale bar = 300 μm. (e) Representative whole-well scans (brightfield, grayscale; GFP, green) demonstrating suppression of RSV-GFP infection at 7 dpi with 6.6 μM RDV. Scale bar = 10 mm. (f) Pre-migration viral load in apical supernatants (copies/μl). RSV604 and remdesivir (RDV) significantly reduce viral load compared to RSV alone (p < 0.05). (g) Pre-migration apical cytokine levels (log2 fold change from mock) for IL-6, IL-8, and IP-10. RSV infection significantly increases cytokines; RSV604 and RDV further enhance these levels (p < 0.05). Error bars represent standard deviation; ns = not significant; *p < 0.05, **p < 0.01

Ciliated AECs (Fig. 6c) were infected with GFP-tagged RSV in the presence and absence of RSV604 or RDV. In untreated controls, viral propagation was evident with increasing numbers of GFP-positive cells over a 7-day period, reflecting active infection and spread. In contrast, samples treated with RSV604 showed no detectable RSV-GFP infection throughout the entire time course, indicating that RSV604 effectively inhibits RSV replication and spread in this model (Fig. 6d). Similarly, ALI cultures treated with RDV showed complete absence of GFP expression compared to untreated controls at 7 days post-infection, confirming effective inhibition of RSV replication in this model (Fig 6e).

Apical supernatants collected before neutrophil introduction revealed a significant reduction in viral load following RDV (1.2 × 10D ± 5.3 × 10D copies/μl) and RSV604 (1.4 × 10D ± 5.0 × 10D copies/μl) treatment compared to untreated controls (3.8 × 10D ± 6.7 × 10D copies/μl, p<0.05, n = 6) (Fig. 6f). These measurements, taken 24 hours post-infection, should be interpreted with caution since plaque assays at this time point were below the detection limit (data not shown), suggesting that the results may not reflect infectious viral load. Meanwhile, pre-treatment with RDV and RSV604 resulted in a significant increase in IL-6 and IP-10 secretion relative to drug treated, uninfected controls (Fig. 6g).

### The Antiviral RSV604 Reduces Neutrophil-Mediated Inflammation During RSV Infection

Finally, we aimed to assess whether RDV or RSV604 treatment could specifically reduce the expression of key neutrophil degranulation markers, MPO and NE, which were significantly elevated in both *in vitro* and *in vivo* models of RSV infection.

Notably, RSV604, but not RDV, led to higher numbers of viable neutrophils recovered from the apical side (Fig. 7a**, Supplementary Fig.8a&b**) and was associated with a lower expression of nearly all degranulation markers on the basolateral side (Fig. 7b). The radar plot demonstrates that RSV604 (pink) markedly shifts neutrophil marker expression, particularly NE and MPO, across all compartments, aligning more closely with mock (grey) conditions (Fig. 7c**, Supplementary Fig.8c**). This suggests RSV604’s potential to effectively modulate the neutrophil response to RSV infection. In contrast, RDV (purple) shows weaker modulation, with marker expression patterns more closely resembling those of the RSV control (blue) or greater (Fig. 7d).

**Figure 7.**
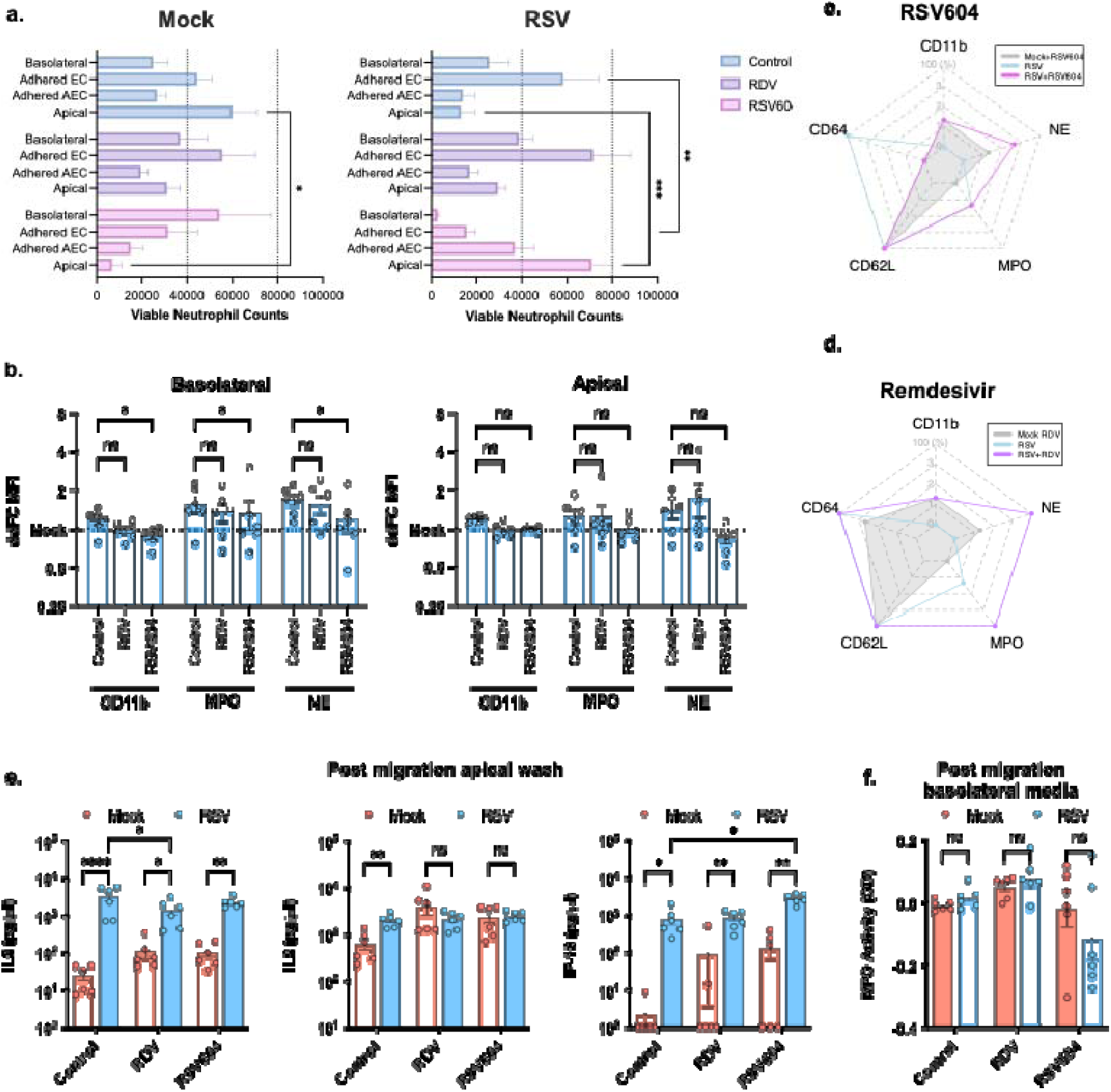
Effects of Antiviral Treatments on Neutrophil Degranulation using in vitro RSV Models. **(a)** Total viable neutrophil counts across compartments. No significant differences (ns) in total neutrophil counts across treatments; RSV604 leads to increased migrated neutrophils. **(b)** Neutrophil marker expression (ΔΔFC of MFI relative to medium-only and mock controls) across compartments and treatments. **(c-d)** Radar plots of neutrophil activation marker quartile scores across compartments (basolateral, adhered epithelial, adhered endothelial, migrated neutrophils). RSV604 reduces MPO and NE activation, especially in migrated neutrophils. **(e)** Post-migration apical cytokine levels of IL-6, IL-8, and IP-10 compared to RSV alone (p < 0.05). Error bars represent standard error; ns = not significant; *p < 0.05, **p < 0.01. **(f)** MPO enzymatic activity in basolateral supernatants. RSV alone increases MPO activity (p < 0.05 vs. mock), while RSV604 and RDV display the same activity levels.

These changes in neutrophil degranulation marker expression corresponded to minor alterations in inflammatory cytokine levels following neutrophil migration. Neutrophil migration following RSV infection and RDV treatment significantly (p<0.05) reduced IL-6 secretions, while RSV604 led to further increase in IP-10 levels from untreated, infected control (Fig. 7e). Treatment with RSV604 showed a trend towards a reduction in MPO enzymatic activity in the basolateral compartment, consistent with the lower MPO expression observed on basolateral neutrophils, although this reduction did not reach statistical significance (Fig. 7f). These findings highlight the potential of antiviral treatments to attenuate neutrophil degranulation during RSV infection.

## Discussion

In this study, we aimed to develop an *in vitro* model that recapitulates key clinical outcomes in infants with RSV bronchiolitis to help accelerate the discovery of effective therapeutics. A key finding was that peripheral blood neutrophils from RSV-infected infants had elevated levels of MPO and NE. Interestingly, these infants exhibited a lower overall neutrophil count in circulation, likely reflecting increased recruitment to the lungs, a well-documented feature of RSV bronchiolitis (8, 10, 31, 32). Elevated MPO and NE levels have also been observed in bronchoalveolar and nasopharyngeal lavage fluid samples from RSV-infected infants (13, 14), and neutrophils recovered from infection sites display activation markers such as increased CD11b and CD64, and reduced CD62L (33–36).

To establish our clinically relevant *in vitro* model, we first investigated whether epithelial age was a principal driver of neutrophil phenotype. Our previous work has shown significant baseline age⍰dependent differences in paediatric versus adult airway epithelium (37), suggesting that epithelial age could be a key determinant of neutrophil behaviour. Here we showed that the paediatric epithelium drives both greater neutrophil migration and a selectively higher MPO and NE signal in basolateral compartment (which models the systemic circulation), closely mirroring the phenotype observed in RSV⍰infected infants.

Building on this, we then established a more physiologically relevant airway microenvironment by incorporating a vascular endothelial layer to simulate neutrophil extravasation during RSV infection (21). As expected for a barrier⍰forming compartment, endothelial cells reduced dextran flux but did not alter RSV replication (viral RNA levels) in the epithelium. RSV exposure induced robust production of IL⍰6, IL⍰8, and IP⍰10 across models, consistent previous *in vitro* studies (11, 38) and with clinical profiles showing elevated IL⍰6, IL⍰8, and IP⍰10 in nasopharyngeal and bronchial aspirates from infants with severe RSV bronchiolitis (39–41). Interestingly, cytokine levels were lower in the EC⍰containing model, suggesting that endothelial cells may modulate early epithelial inflammatory signalling, a finding that contrasts with influenza A virus, where ECs have been shown to amplify epithelial cytokine production (42).

Using this model, we found that neutrophil trans-epithelial migration increased markedly following RSV infection, likely driven by elevated IL⍰8 and IP⍰10. In the absence of endothelial cells, most neutrophils remained adherent to RSV⍰infected airway epithelium, whereas the EC⍰containing system supported a substantially greater number of fully migrated, detached neutrophils. This is consistent with previous work (11, 18), and suggests that endothelial cells influence neutrophil behaviour, with migrated neutrophils potentially contributing more extensively to RSV⍰associated pathology. We found significant differences in basolateral neutrophil expression of MPO and NE, but no other significant differences between RSV and mock conditions were detected. These findings are consistent with the dominant MPO/NE⍰driven signature reported previously in an adult *in vitro* RSV migration system (12). However, that study showed a stepwise increase in CD11b, NE, and MPO with migration, with the highest levels in apical neutrophils. These differences are likely due to differences in epithelial age in the two systems, highlighting the importance of epithelial-neutrophil interactions is determining phenotype. While not the aim of this current study, a logical next step would be to define the adhesion pathways underpinning these age⍰dependent effects. Prior work has shown that LFA⍰1:ICAM⍰1 interactions are essential for neutrophil transepithelial migration in RSV infection (18), blocking this interaction could help elucidate a mechanistic link between epithelial age and neutrophil response.

Using quartile⍰based analyses across all markers we found that migrated neutrophils displayed a distinct activated phenotype (elevated CD11b, CD64, MPO, and NE, with reduced CD62L), mirroring profiles seen in bronchoalveolar lavage fluid and peripheral blood neutrophils from RSV-infected infants (33–36, 43, 44). A key finding was that this basolateral neutrophil phenotype appeared to depend on direct contact with RSV-infected epithelial cells rather than soluble factors alone. Notably, basolateral neutrophils that were prevented from migrating did not acquire this phenotype, supporting the hypothesis that trans⍰epithelial migration is necessary for full activation. These findings raise the possibility that basolateral neutrophils may re⍰enter the circulation after activation, with migration across endothelial and RSV⍰infected epithelial layers priming them for degranulation and potentially contributing to systemic inflammation by transporting viral components beyond the lung (11, 45, 46).

Finally, we used this model to investigate the effects of two well-characterized RSV inhibitors, RDV and RSV604, which target distinct stages of RSV replication (28, 47). We found that both significantly reduced viral load and GFP fluorescence in infected AECs, consistent with previous studies (28, 48). However, the reduction in IL-6 levels with RDV, observed only after neutrophil migration, suggests that the presence of neutrophils, possibly in conjunction with reduced viral antigen load, may contribute to a regulatory feedback mechanism that reduces inflammation thereby improving disease outcomes.

Interestingly, RSV604 increased the number of viable neutrophils migrating across infected co-cultures. Neutrophils play a crucial role in viral clearance, but excessive recruitment and degranulation can cause tissue damage. However, our data indicate that neutrophils migrating under RSV604 treatment were less activated, with reduced expression of CD11b, MPO and NE. This suggests that RSV604 may promote the recruitment of less activated, “naive” neutrophils, which could mitigate excessive inflammation. Further research is needed to explore how RSV604 influences these pathways and whether it has the potential to modulate neutrophils *in vivo*. A working hypothesis is shown in Fig. 8.

**Figure 8.**
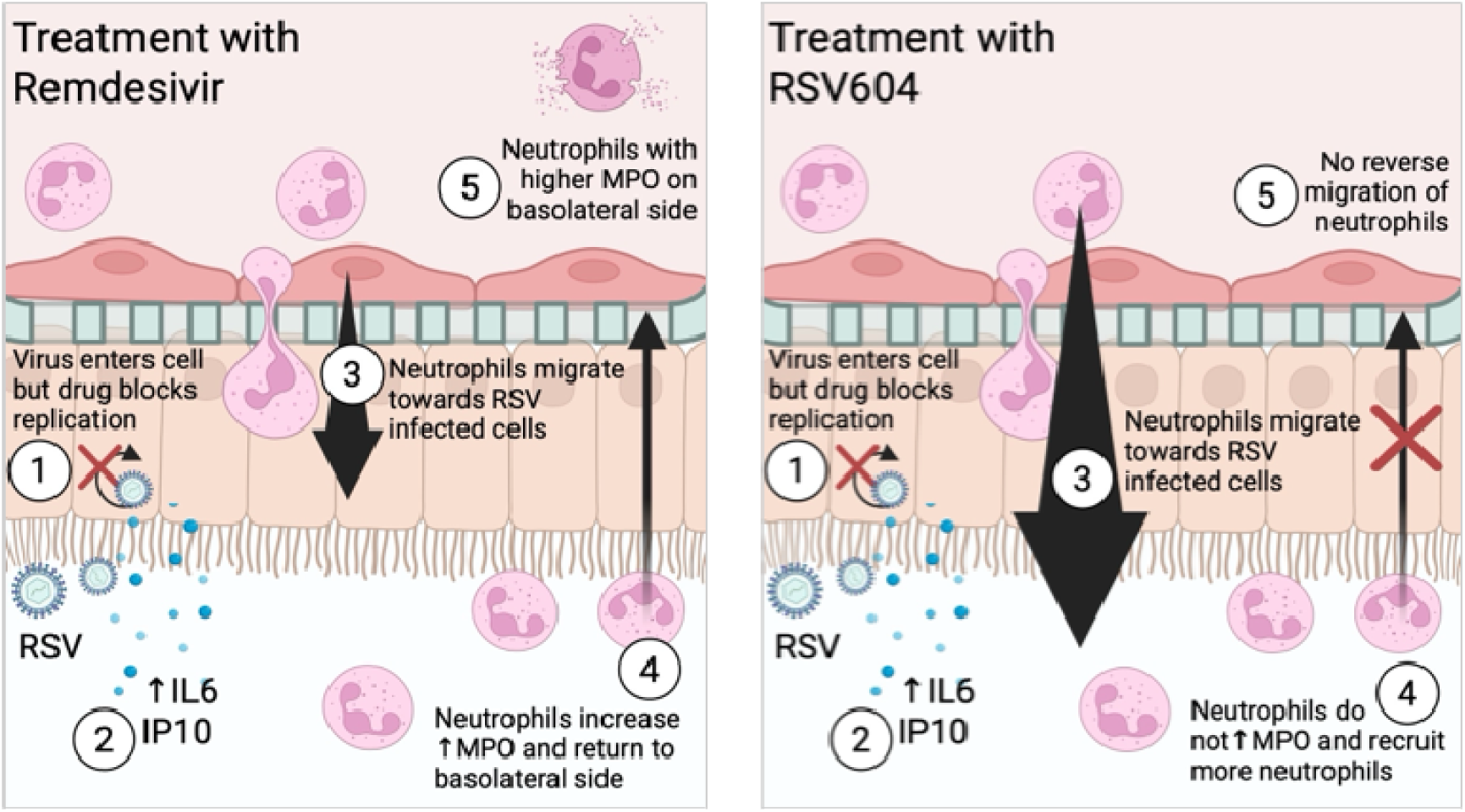
Proposed model of neutrophil migration and activation during RSV infection with and without RSV604 treatment. Schematic diagrams depict epithelial infection, cytokine secretion, and neutrophil recruitment under remdesivir (left) and RSV604 (right) treatment conditions. In the remdesivir model, infected epithelial cells release IL/JI6 and IP/JI10, leading to neutrophil activation, increased MPO and NE expression, and basolateral migration with elevated degranulation. In contrast, RSV604 reduces epithelial cell infection and inflammatory signalling, resulting in neutrophil migration with lower CD11b, MPO and NE expression. Together, these diagrams illustrate a potential mechanism by which RSV604 promotes the recruitment of less/JIactivated neutrophils and limits tissue/JIdamaging inflammation.

Limitations of this study include the use of paediatric neutrophils for *in vivo* analysis and adult neutrophils for *in vitro* experiments. This was principally due to logistical challenges in obtaining healthy paediatric blood samples on the morning of the migration assay. Although direct comparison between healthy paediatric and adult donors revealed only differences in the resting state of CD64, other age-related differences in neutrophil responsiveness or priming potential may still influence the observed outcomes. Additionally, the *in vitro* model, while physiologically relevant, does not fully recapitulate the complexity of the *in vivo* airway environment, including interactions with other immune cell types and systemic factors.

## Conclusion

Our study identifies myeloperoxidase (MPO) as a key marker of neutrophil activation and a potential systemic indicator of disease severity in RSV infection. Using a clinically informed *in vitro* model, we showed that neutrophil activation is primarily driven by direct contact with RSV-infected epithelial cells, resulting in increased expression of CD11b, MPO, and NE, features associated with severe disease. MPO emerged as the most significantly modulated marker, confirmed by both expression and enzymatic activity assays. Although both RDV and RSV604 reduced MPO levels compared to RSV alone, neither restored them to baseline, indicating only partial suppression of degranulation. Notably, only RSV604 effectively reduced MPO expression, suggesting its potential to mitigate neutrophil-mediated tissue damage. Furthermore, high MPO expression in basolateral neutrophils points to a possible role in systemic inflammation. These results suggest that neutrophil MPO expression should be considered a relevant biomarker in the evaluation and screening of antiviral therapies for RSV.

## Acknowledgements

This study was funded by Animal Free Research UK (AFR19-20274) and was carried out in accordance with the terms and conditions as stated in document number GH2019-TC/v1.1/27.01.2020. We would like to thank the members of the research nursing team at GOSH, including Lauren O’Neill, Holly Bellfield, Shannon McCarty, Kate Plant and Sarah Benkenstein, who helped with patient screening and sample collection. This work was supported by the NIHR Great Ormond Street Hospital Biomedical Research Centre. We are grateful to Prof. Cecilia Johannssen for her guidance during initial project development and for review of the manuscript. The views expressed are those of the author(s) and not necessarily those of the NHS, the NIHR or the Department of Health. C.M.S also acknowledges funding support from UKRI/ BBSRC (BB/V006738/1).

## Author contributions

M.P. designed the study, conducted experiments, analysed data, and co-wrote the manuscript. T.M. and E.J.R. conducted experiments, analysed data, and co-wrote the manuscript. W.C. conducted experiments and analysed data. L.B. assisted T.M. with transcriptomic analysis. S.E. and A.I.J. contributed to study design, conducted experiments and provided access to reagents. S.E. also assisted in reviewing the manuscript. T.B. and P.D.C. provided access to reagents. C.M.S. and S.R. provided support through ethics and patient recruitment. R.L.S. and C.M.S. conceived the study and oversaw the funding application. J.L.R. and G.P. contributed to data analysis (quartile scoring). M.C-B contributed to statistical analyses. M.W contributed to study design. C.M.S. contributed to study design and co-wrote the manuscript. M.P., R.E.H., and C.M.S. oversaw data analysis, interpretation, and the manuscript write-up.

## Code availability

Custom code for the analysis performed in this study will be made publicly available via GitHub at https://github.com/smithlab-code.

---

**Supplementary Figure 1.**
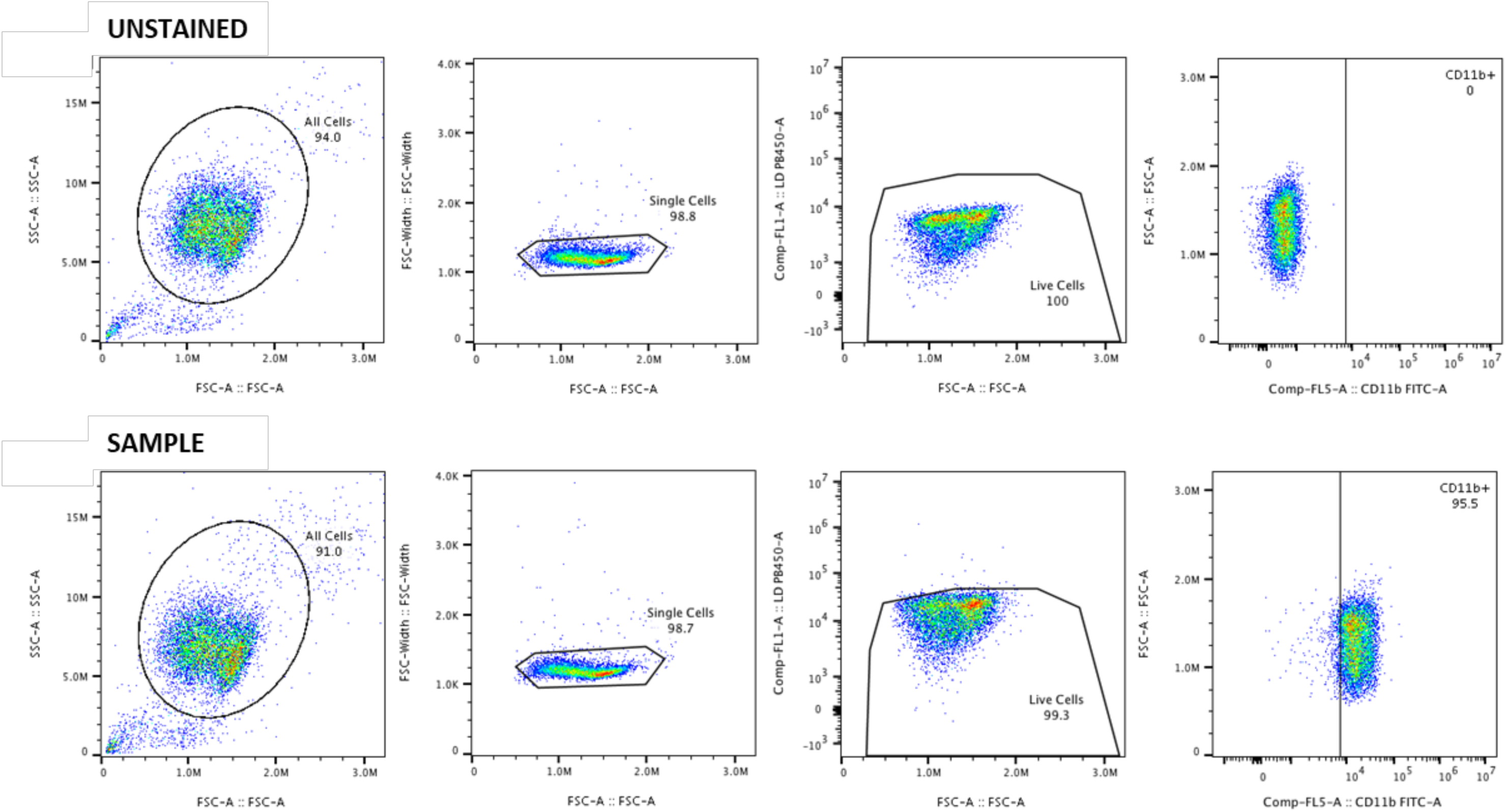

**Supplementary Figure 2.**
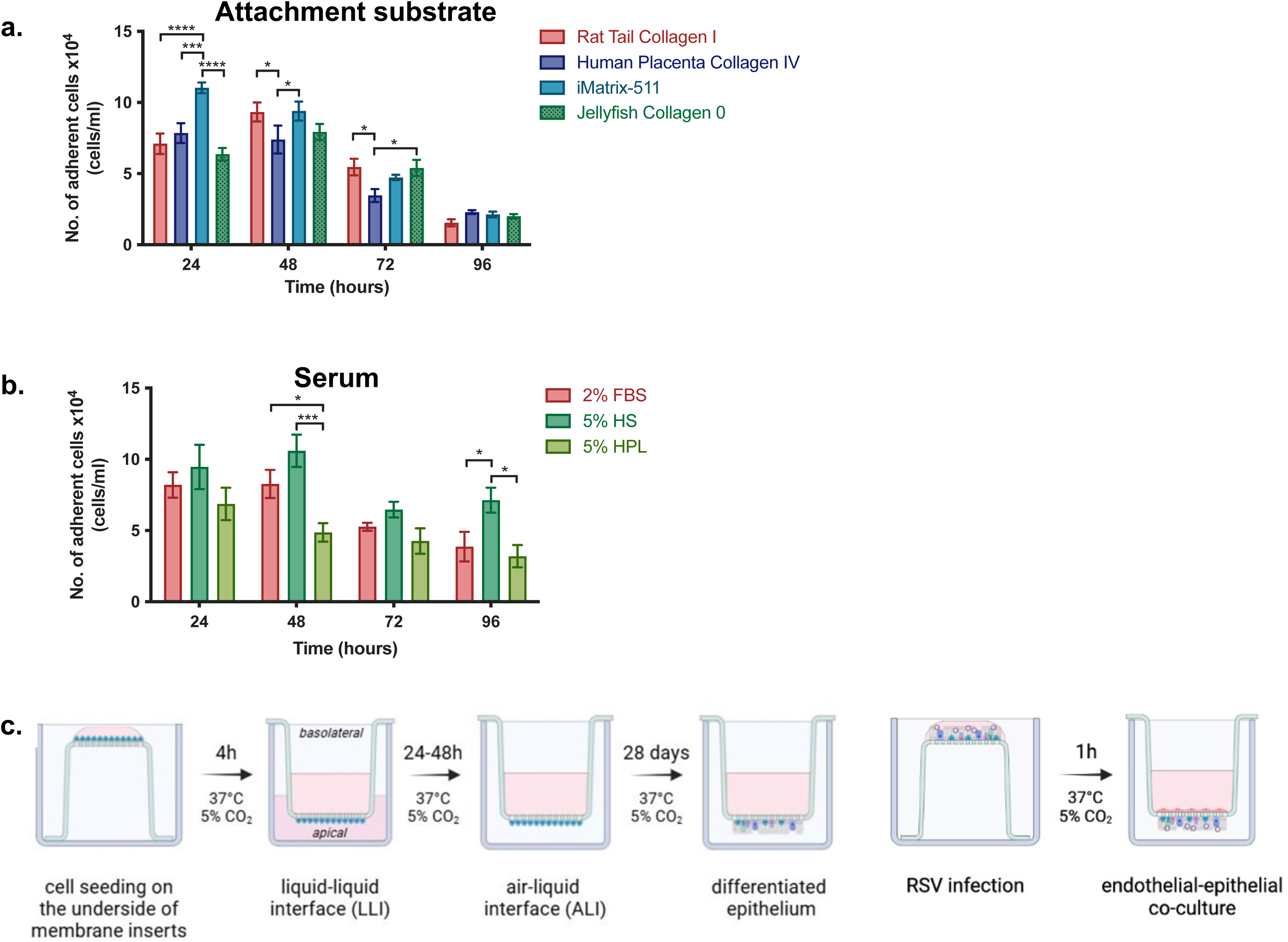

**Supplementary Figure 3.**
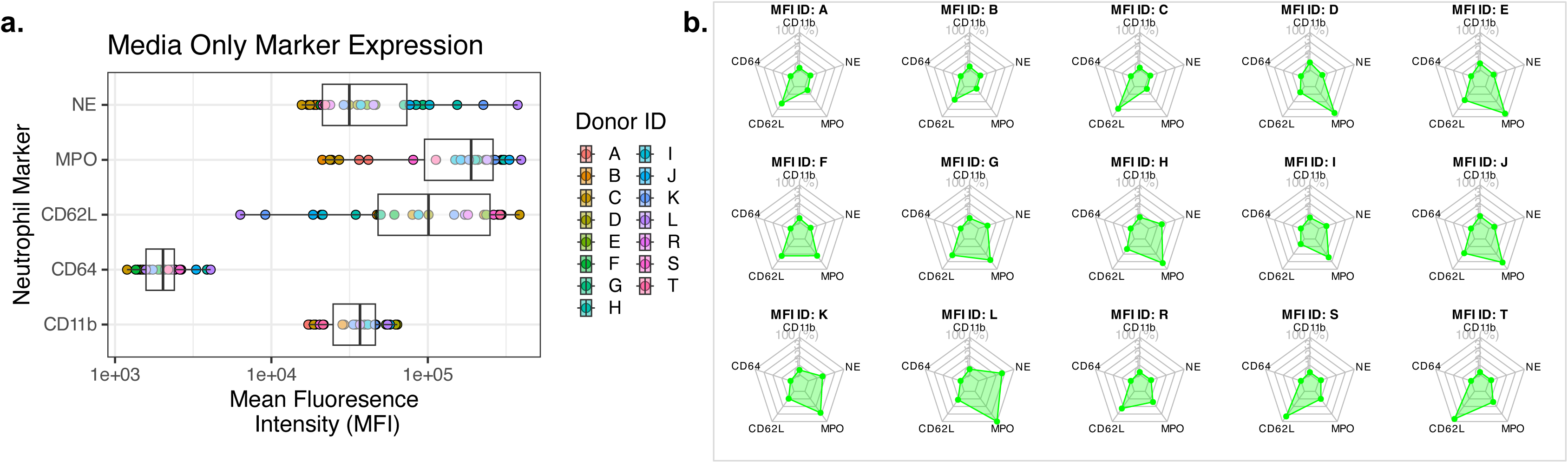

**Supplementary Figure 4.**
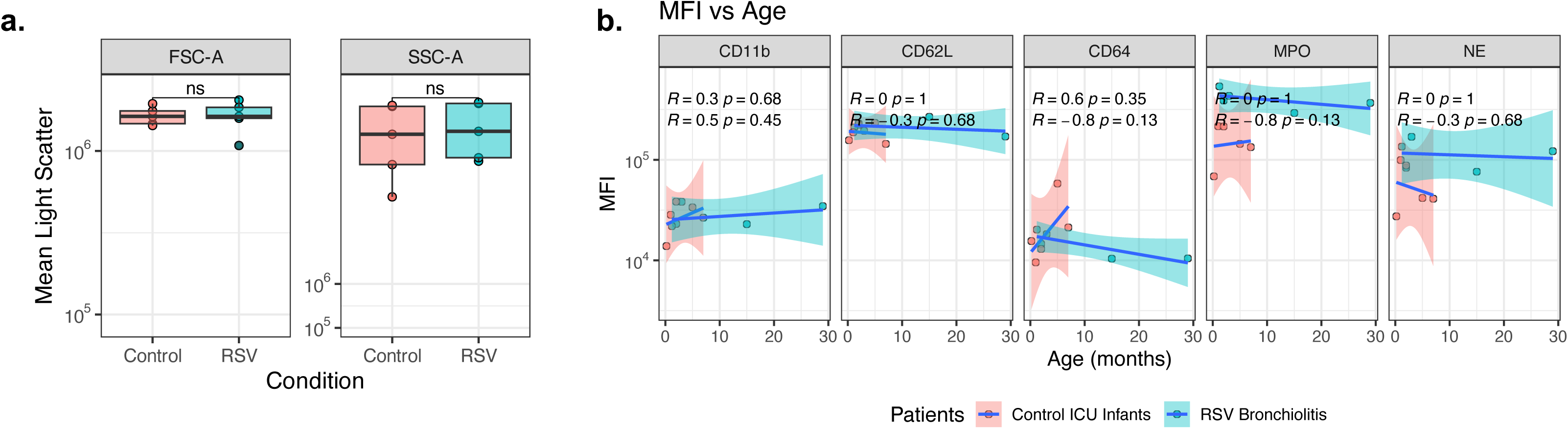

**Supplementary Figure 5.**
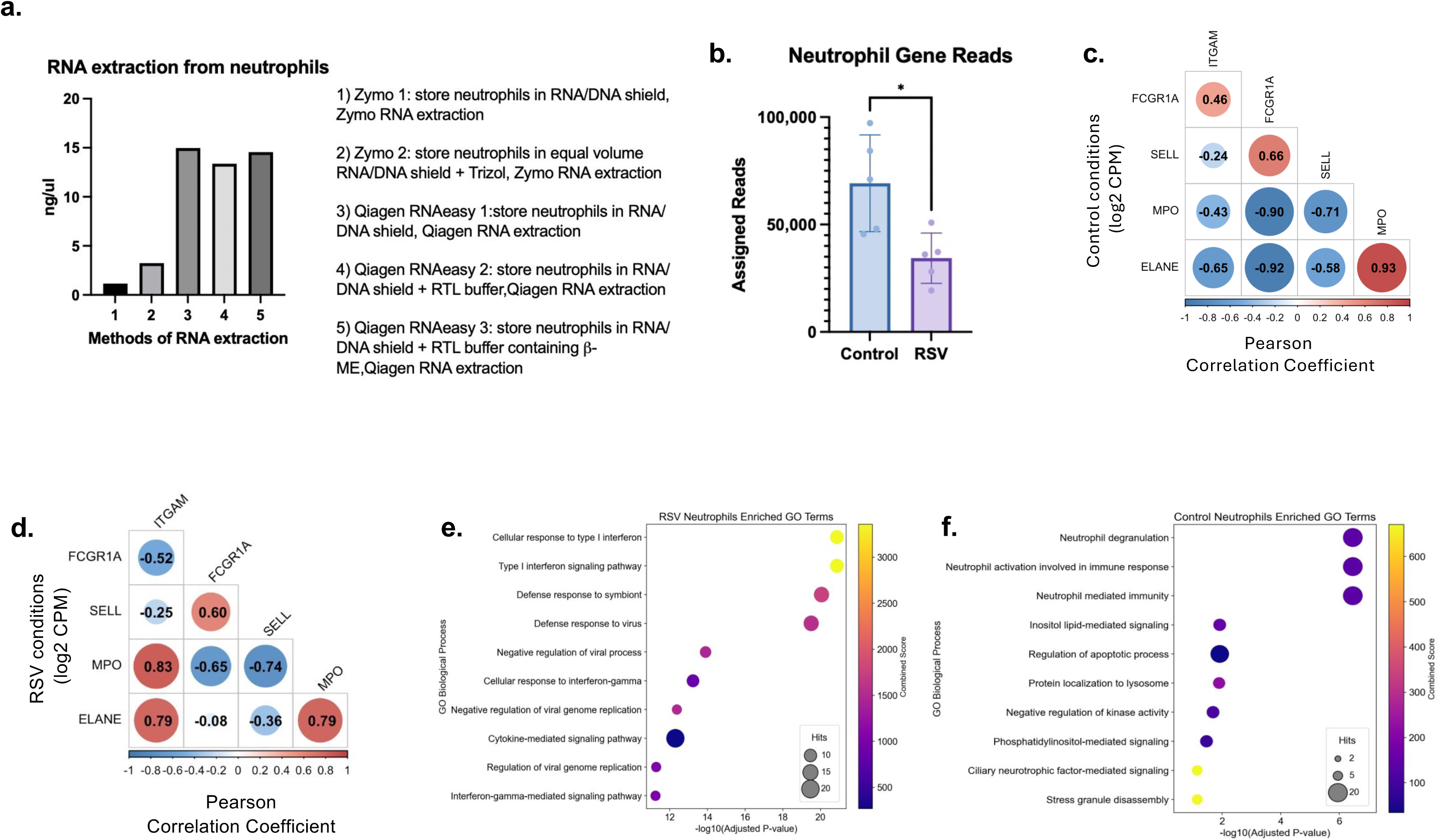

**Supplementary Figure 6.**
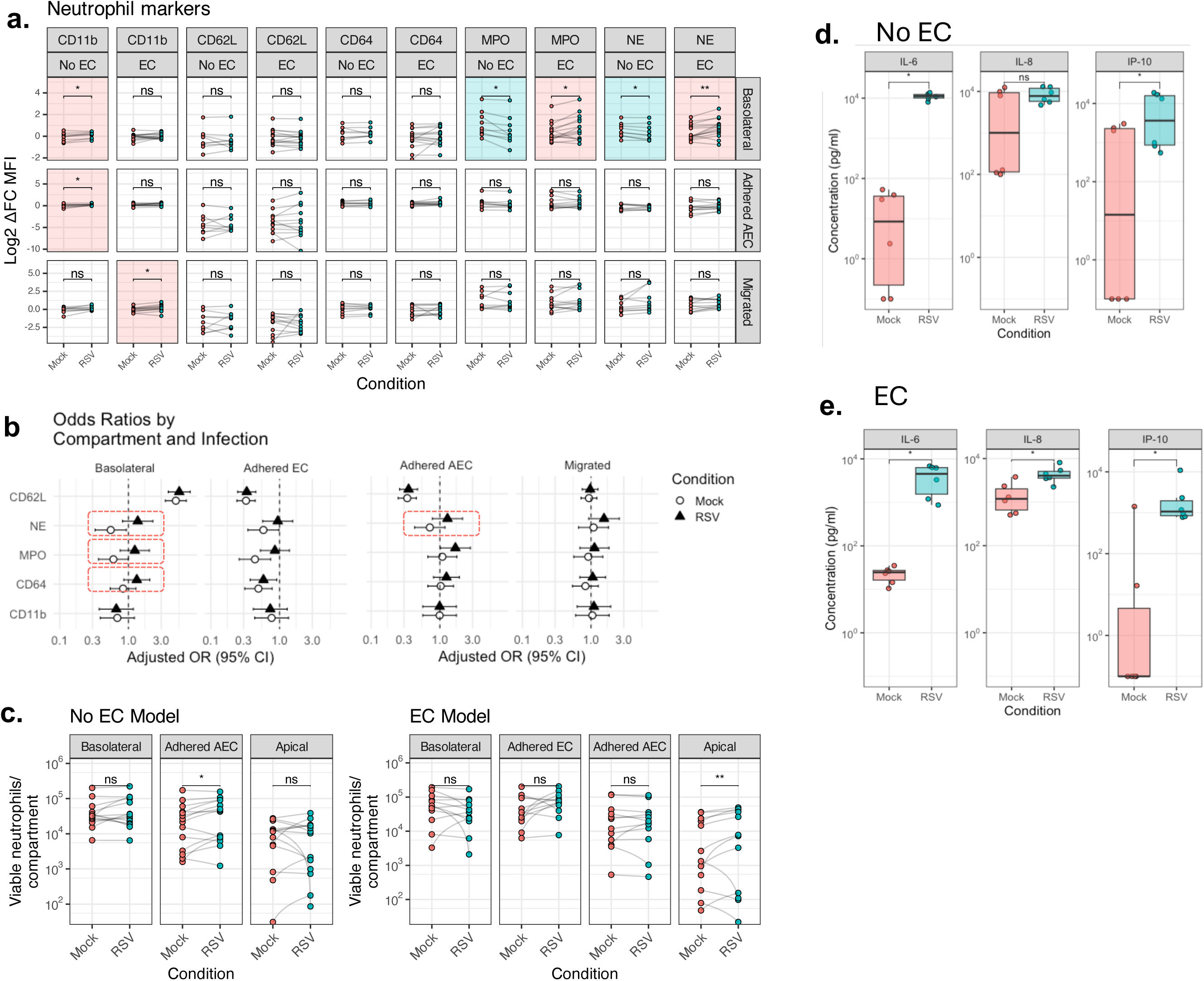

**Supplementary Figure 7.**
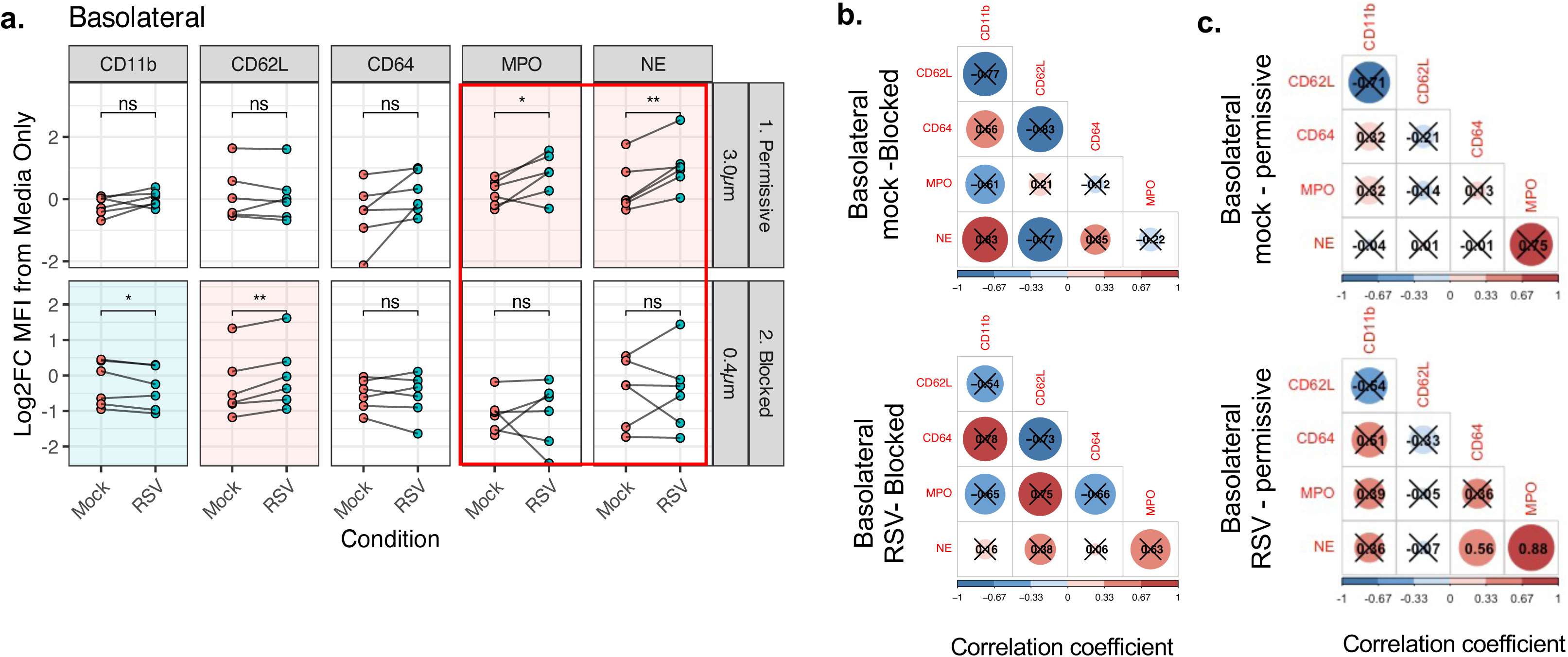

**Supplementary Figure 8.**
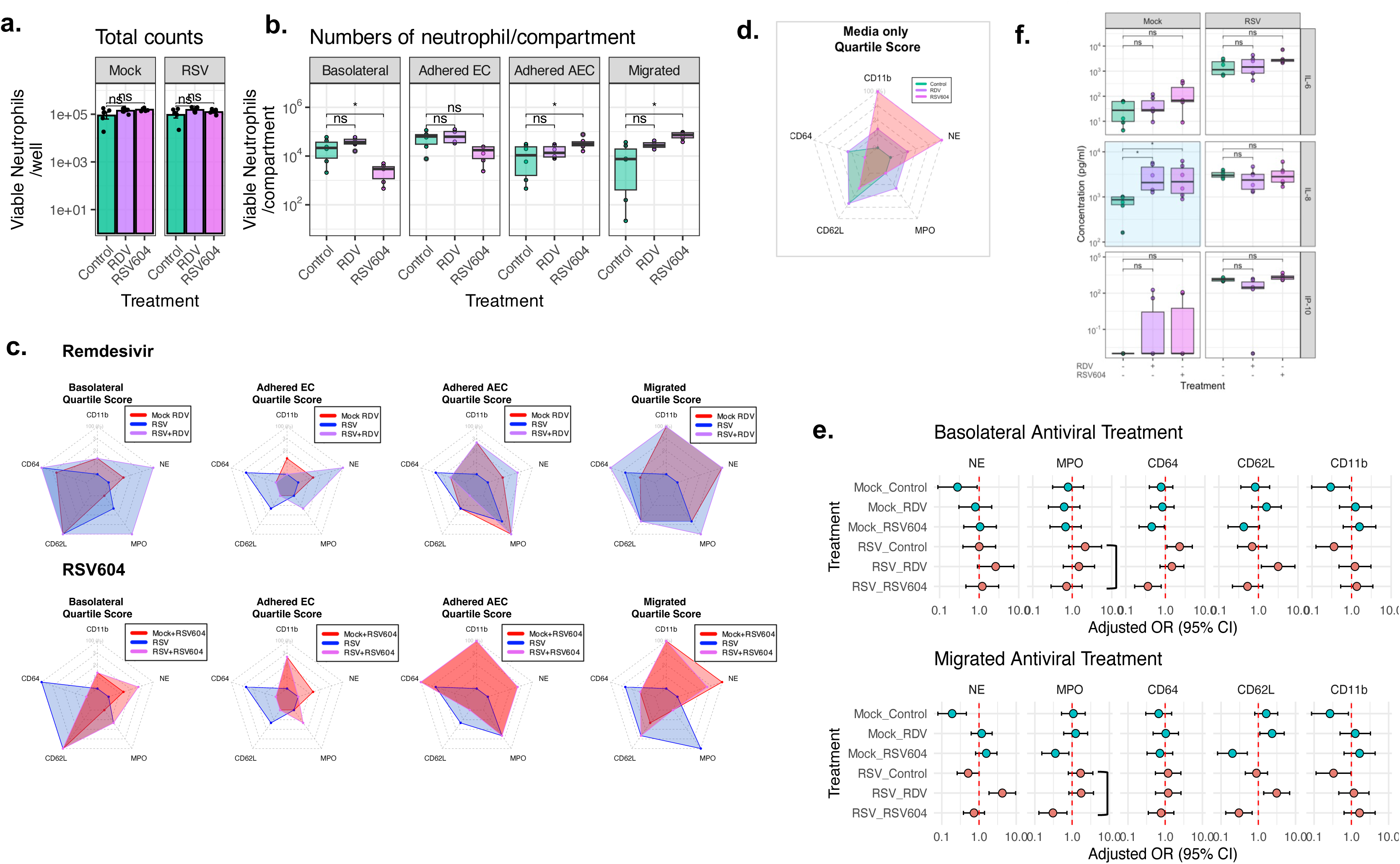

## Notes

### Competing Interest Statement

The authors have declared no competing interest.

### Summary of Updates

The clinical dataset has been re-analysed using age-adjusted statistical models to clarify associations between RSV infection and neutrophil activation markers; corresponding updates have been made in the Results, Figure 1f, and Supplementary Figure 4b. We added new comparisons of adult and paediatric neutrophils, clarifying age-related differences and revising text to moderate claims regarding neonatal neutrophils. Mechanistic insight has been expanded through the addition of infant neutrophil transcriptomic profiling aligned with in‑vitro phenotypes (new panels in Figure 2) and through new time-lapse imaging of neutrophil migration (Figure 3d-f). We also incorporated experiments separating soluble factor-dependent from contact-dependent effects using permeable versus non-permeable inserts (Figure 5). The role of endothelial cells has been clarified, with new text emphasising their contribution to barrier context rather than epithelial infection. Corresponding updates were made to Figure 4 and the Methods. Across all figures, donor numbers, replication structure, and assay schematics have been added or clarified. Flow cytometry reporting has been improved with representative plots incorporated into main figures and full gating strategies in Supplementary Figure 1.

