## Supplementary Figure Legends for "Neutrophil myeloperoxidase as a functional biomarker for RSV severity: implications for *in vitro* therapeutic screening"

**Supplementary Figure 1.** Overview of flow cytometry gating strategy for neutrophils.

**Supplementary Figure 2.** Impact of Attachment Substrate and Serum on Endothelial Cell Culture. (a) Comparison of cell attachment across different substrate conditions. (b) Effect of serum presence on cell recovery.

**Supplementary Figure 3.** Donor Variability in Neutrophil Marker Expression and Cytokine Production in Response to RSV. (a) Mean fluorescence intensity (MFI) of neutrophil markers (NE, MPO, CD62L, CD64, CD11b) for individual donors under media‑only conditions, demonstrating substantial inter‑donor variation. (b) Radar plots displaying donor‑specific neutrophil activation profiles (donor IDs A–T), highlighting heterogeneity in marker expression.

**Supplementary Figure 4.** Additional Analyses of Infant Sample Variability. (a) Comparison of forward scatter (FSC‑A) and side scatter (SSC‑A) between infant control and infant RSV bronchiolitis samples, b) Correlation plots showing the relationship between infant age and mean fluorescence intensity (MFI) of neutrophil activation markers, showing no age‑associated trends within infant samples.

**Supplementary Figure 5.** Transcriptomics Quality Control. (a) Comparison of five different neutrophil RNA extraction methods. (b) Assigned sequencing reads for purified neutrophils from control and RSV‑infected infants. Bars show mean ± SD, demonstrating reduced transcript capture in RSV samples. (c) Pearson correlation matrix of key neutrophil genes under control conditions (log₂ CPM). Higher positive correlations are shown in red, negative correlations in blue. (d) Pearson correlation matrix of the same genes under RSV‑infected conditions (log₂ CPM). (e) Gene ontology (GO) enrichment analysis of pathways upregulated in RSV neutrophils. Dot plot displays the top enriched GO biological processes, with dot size indicating number of genes per pathway and colour indicating –log₁₀ adjusted p‑value. (f) GO enrichment analysis of pathways associated with control neutrophils.

**Supplementary Figure 6.** Impact of Neutrophil Migration on Viral Load, Cytokine Production, and Neutrophil Distribution in Models with and without Endothelial Cells (EC). (a–b) Additional data visualisation outputs showing differences in infection, compartment or immune response. (c) Distribution of neutrophils across model compartments (basolateral, adhered AEC, apical) in EC and No‑EC systems under RSV and mock conditions; no significant differences observed. (d) Viral load and cytokine concentrations (IL‑6, IL‑8, IP‑10) in “No EC” epithelial‑only models. (e) Viral load and cytokines in models incorporating endothelial cells. RSV significantly increases viral replication (p < 0.01) and cytokine production (p < 0.05) relative to mock. Error bars = SD; ns = not significant; *p < 0.05, **p < 0.01.

**Supplementary Figure 7.** Correlation of Neutrophil Marker Expression in the Basolateral Blocked Condition. (a) Paired analysis of neutrophil marker expression across experimental conditions, illustrating comparative changes in activation marker levels. (b–c) Correlation matrices for neutrophil activation markers (CD11b, CD62L, CD64, MPO, NE) across different experimental conditions (mock‑blocked, mock‑permissive, RSV‑permissive, RSV‑blocked). Positive correlations are shown in red and negative in blue; only statistically significant correlations are displayed. Heatmaps illustrate how marker relationships vary depending on epithelial/endothelial permissiveness and infection status.

**Supplementary Figure 8.** Impact of Antiviral Treatments on Cytokine Production and Neutrophil Activation in RSV Models. (a) Total counts of neutrophils recovered from all compartments, demonstrating no loss of cell viability across treatment conditions. (b) Total neutrophil counts separated by compartment (basolateral, adhered, migrated), confirming comparable recovery in each compartment. (c–d) Quartile‑scaled radar plots of mean fluorescence intensity (MFI) for neutrophil activation markers (CD11b, CD62L, CD64, MPO, NE) in basolateral and migrated compartments under RSV604, Remdesivir (RDV), and control conditions. (e) Cytokine concentrations (IL‑6, IL‑8, IP‑10) measured across Mock, RSV, RDV, and RSV604 treatment groups. Error bars = SD; ns = not significant; *p < 0.05, **p < 0.01.
